# Protocol for constructing correlation-based molecular networks from large-scale untargeted metabolomics data

**DOI:** 10.1101/2025.04.26.649581

**Authors:** Huang Lin, Lijun Zhang, Ali Lotfi, Alan Jarmusch, Iris Lee, Adam Kim, James T. Morton, Alexander Aksenov

## Abstract

This protocol describes a computational approach for constructing correlation-based molecular networks from untargeted metabolomics data using MetVAE, a variational autoencoder-based framework. Complementing spectral similarity networks, it captures functional relationships reflected in cross-sample correlations. The workflow imports metabolomics features and sample metadata, adjusts for compositionality, missingness, confounding, and high-dimensionality, estimates sparse metabolite correlations, and exports GraphML files for network visualization. In a hepatocellular carcinoma mouse model, it links lipid classes in high-fat-diet animals, suggesting an endogenous “auto-brewery” route to lipotoxic metabolites.

## Before you begin

Metabolomics is widely used to characterize small-molecule variation associated with disease, treatment, and environmental exposure^1^, but in untargeted mass spectrometry^2–4^ datasets only a minority of detected features can usually be annotated with confidence^5–7^. As a result, biological interpretation often depends on patterns of abundance variation across samples rather than on known metabolite identities alone. Molecular networking based on MS/MS spectral similarity has become an important framework for organizing these data^8–12^, but spectral similarity captures structural relatedness and may miss biologically meaningful associations among structurally dissimilar metabolites that co-vary because of shared sources, pathway context, or coordinated regulation^13–15^.

This protocol describes a computational workflow for constructing correlation-based molecular networks using Metabolomics Variational Autoencoder (MetVAE). The workflow is intended for large-scale LC-MS or GC-MS studies in which relative abundances, missing values, technical confounding, and high dimensionality complicate direct correlation analysis. MetVAE addresses these issues by combining centered log-ratio (CLR)-based preprocessing, Variational Autoencoder (VAE)-based reconstruction, covariate adjustment, and sparse correlation estimation within a single workflow. Although the protocol is motivated by the application to untargeted hepatocellular carcinoma mouse data, the same computational strategy can be adapted to other study-specific or repository-scale metabolomics datasets.

Alternative approaches for sparse correlation or network inference include Pearson correlation, SparCC^16^, SECOM^17^, SPIEC-EASI^18^, and SPRING^19^, each of which addresses only part of issues of the untargeted metabolomics data. In the associated benchmark analyses, we compared these approaches across simulated metabolomics scenarios. Interested readers can find the simulation design and benchmarking results in the Supplementary Information (Figures S1 and S2). With that context in mind, the following practical considerations are important before beginning the workflow.

**CRITICAL:** Untargeted metabolomics data are compositional and often contain extensive missingness. Avoid applying ad hoc pseudo-count, complete-case, or unadjusted pairwise-correlation workflows before this protocol, as these choices can distort correlations and obscure biologically meaningful network structure.

Before beginning the computational workflow, prepare the dataset and analysis plan with the following considerations in mind.

1. Assemble a processed metabolomics feature table and matched sample metadata before beginning the workflow. The input data should already have undergone peak detection, alignment, and feature quantification, and each sample in the abundance matrix must map unambiguously to its metadata record.
2. Define the biological comparison and candidate covariates in advance. Distinguish technical variables, such as batch, instrument platform, acquisition date, and processing run, from biological or sociodemographic/clinical variables, such as diet, disease status, age, sex, and treatment. These variables should be identified before model fitting because the protocol uses covariates to reduce technical bias and clarify downstream correlation estimates.
3. Confirm that the dataset is appropriate for correlation-based network analysis. This approach is most useful when the study contains enough samples to estimate reproducible co-variation patterns across features and when a substantial proportion of metabolites are measured as relative rather than absolute abundances.
4. Review feature sparsity and remove entries that are too rare or too poorly measured for stable downstream modeling. The exact filtering threshold may vary by platform and study size, but features observed in only a very small fraction of samples generally contribute little interpretable network information.
5. Decide whether the protocol will be used primarily for biological discovery, data-structure diagnostics, or both. In the associated work, correlation networks were used to identify coordinated lipid modules, generate hypotheses about shared metabolic origin, reduce technical variation, and flag likely feature-splitting artifacts.
6. Prepare the expected outputs for downstream interpretation. The workflow produces reconstructed feature values, a sparse positive-definite correlation matrix, and a correlation-based network that can be visualized alone or compared with conventional spectral-similarity molecular networks to distinguish functional from structural relationships.

**Note:** If annotated metabolites are available, module-level interpretation is often more direct. If annotations are sparse, the same workflow can still prioritize co-varying unknown features and guide targeted follow-up using spectral libraries, molecular networking, or orthogonal metadata.

## Quickstart: Run all steps using default parameters

### Timing: 3 s

Optional: The full computational workflow is described in detail in the *Step-by-step method details* section. This Quickstart section provides a minimal example for running MetVAE in Python using a small simulated metabolomics dataset with aligned sample metadata.

1. Install the MetVAE package and import the required Python dependencies.

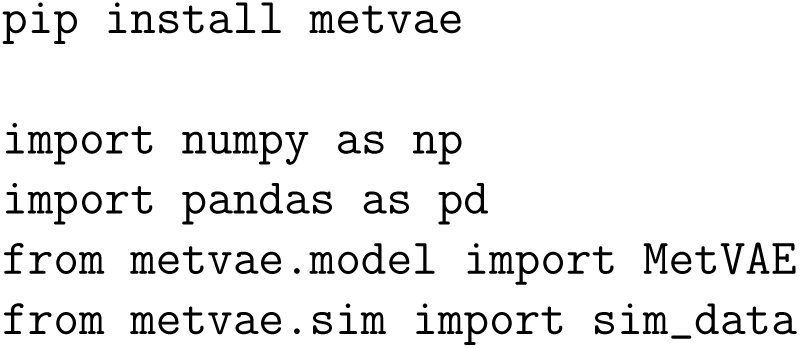
2. Create a small simulated metabolomics dataset and matching sample metadata.
  (a) The feature table contains samples in rows and metabolite features in columns (Figure 1A).
  (b) The metadata table contains one row per sample and aligned sample identifiers (Figure 1B).
  (c) The simulated example includes one continuous covariate (`age`) and one categorical covariate (`batch`).

**Figure 1.**
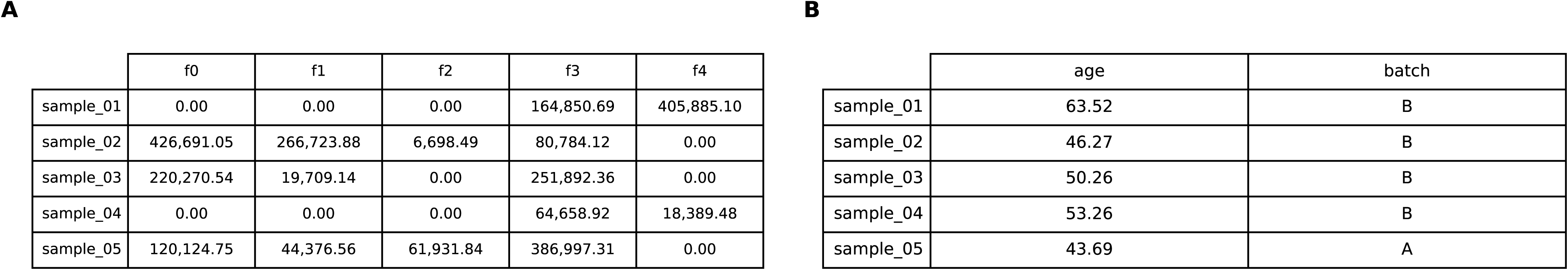
The expected format of the input data. (A) A matrix of metabolite abundances with samples in rows and metabolite features in columns. (B) A metadata table with one row per sample and aligned sample identifiers. In the Quick-start example, the metadata include one continuous covariate (`age`) and one categorical covari-ate (`batch`).

**Note:** This simulated example is intended to illustrate the workflow without requiring study-specific preprocessing.

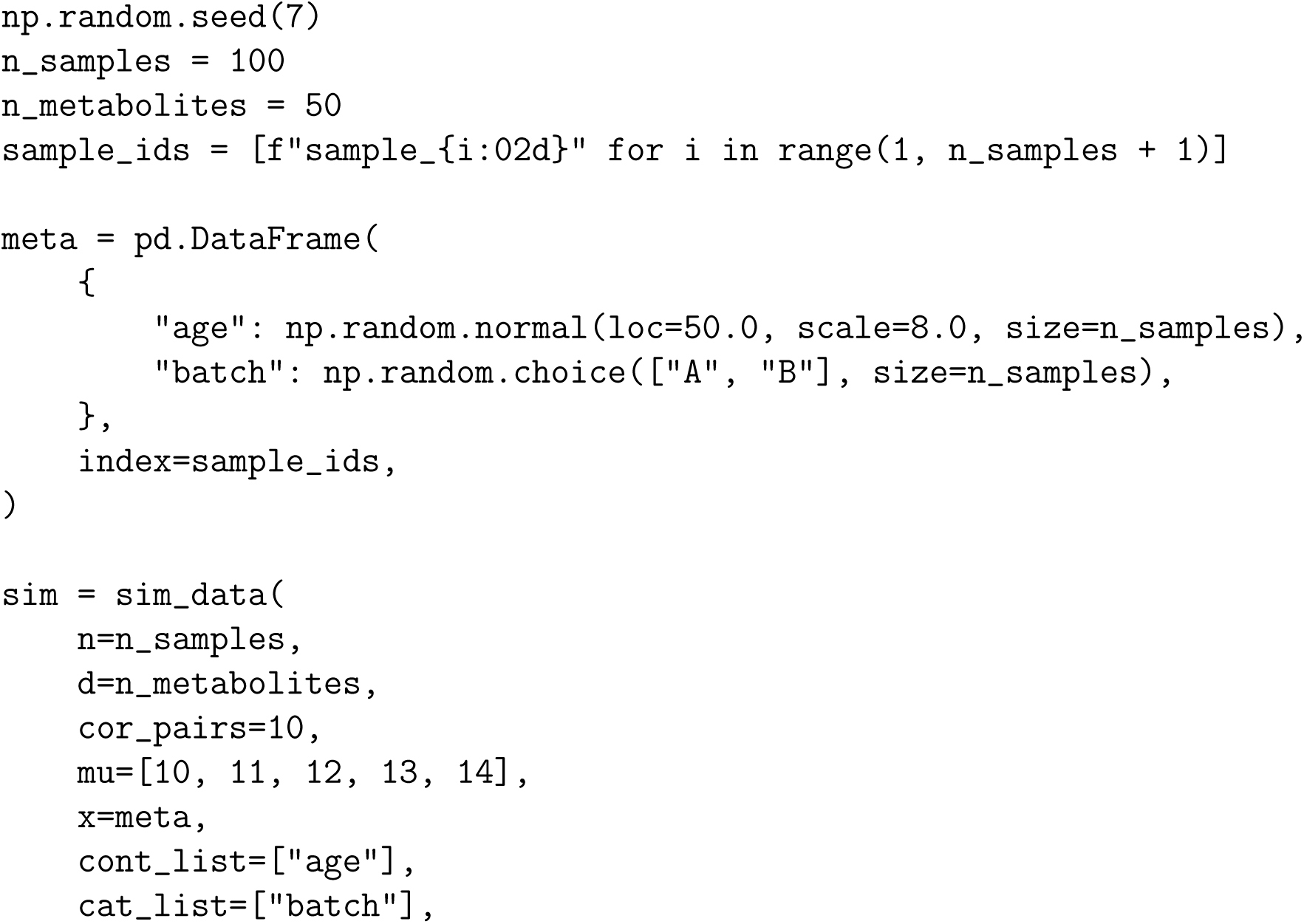
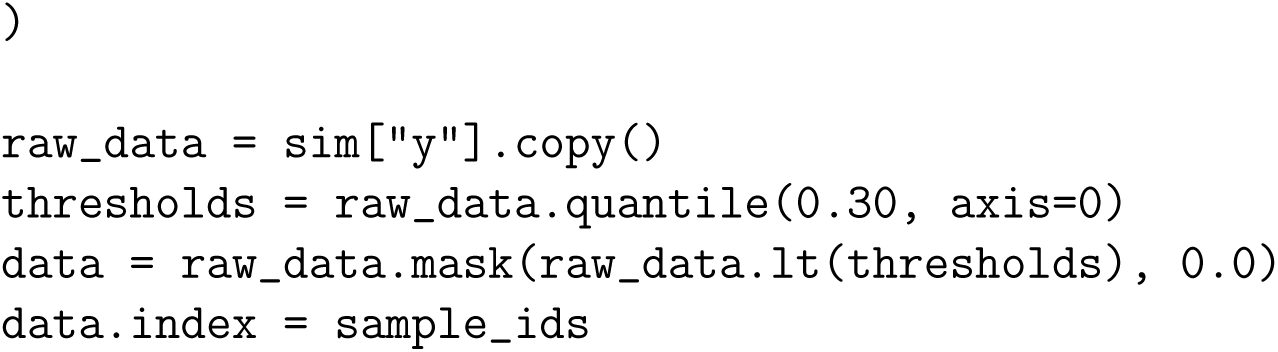

3. Inspect the input structure, initialize MetVAE, and train the model.

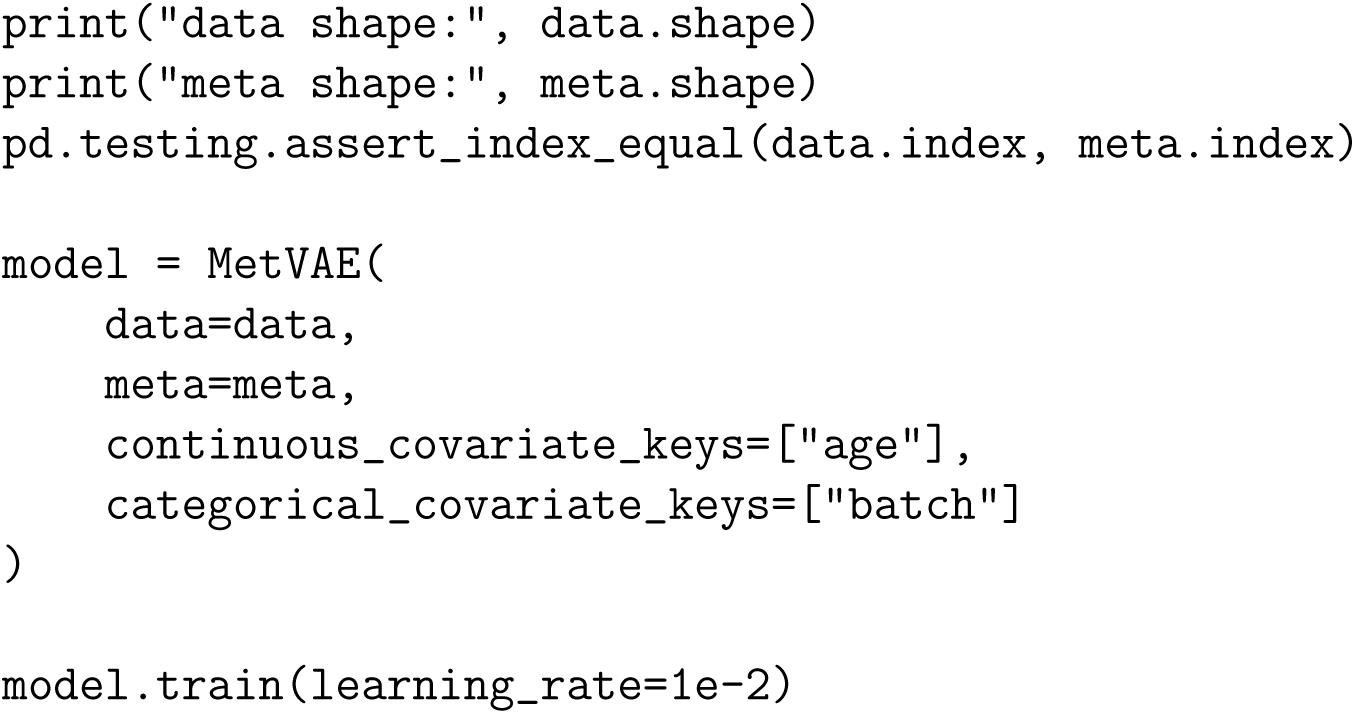

4. Estimate correlations and apply the two built-in sparsification strategies.

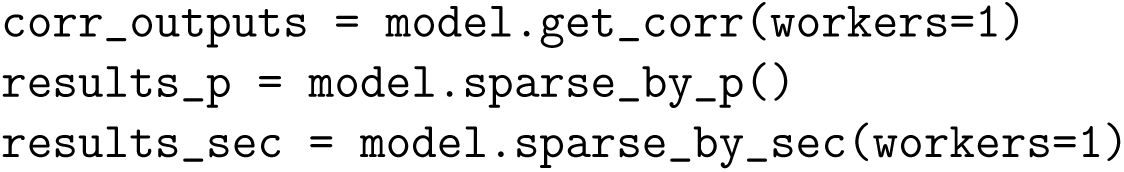

5. Optionally compare both sparsification methods with the known simulated truth.

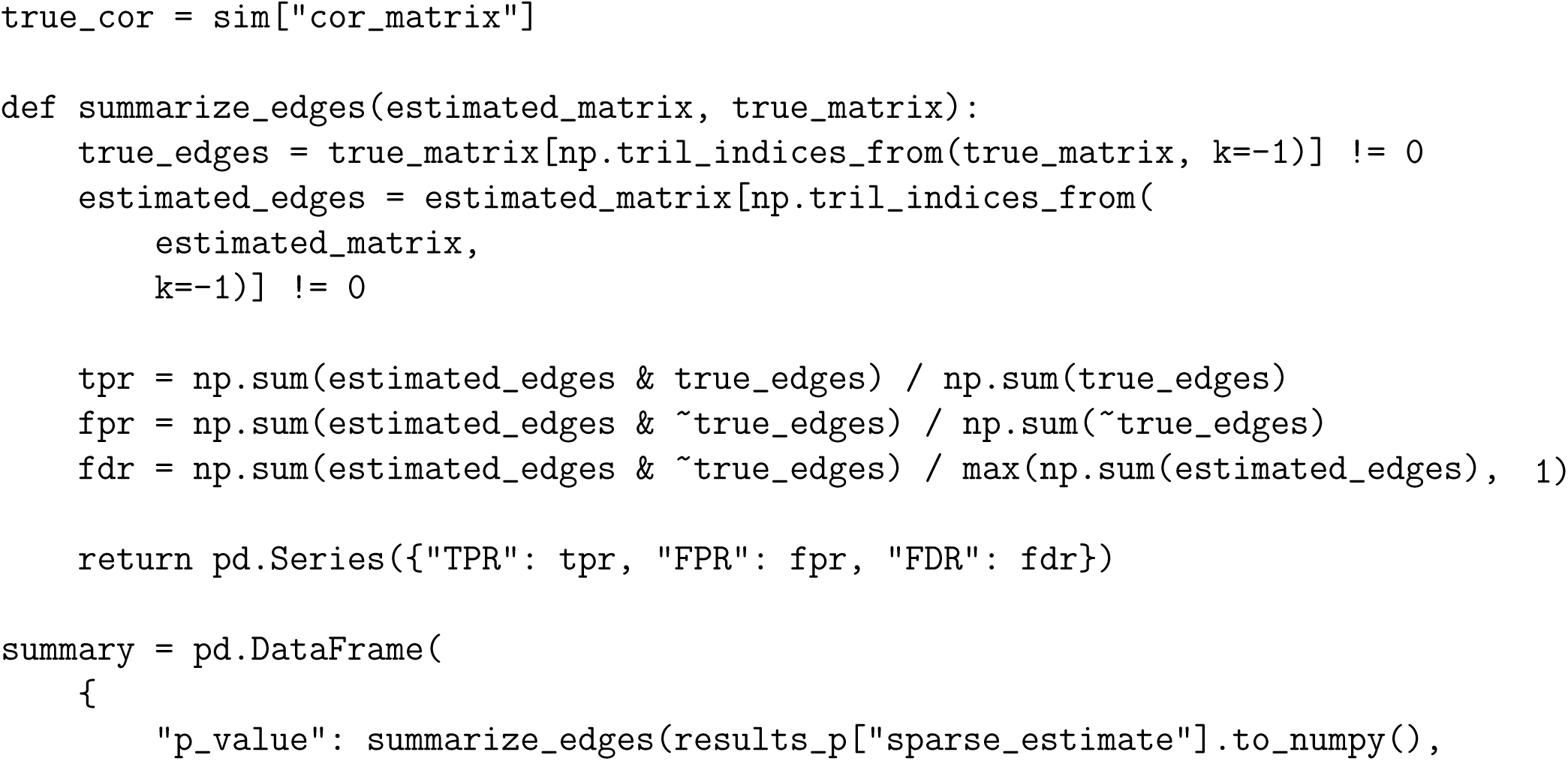
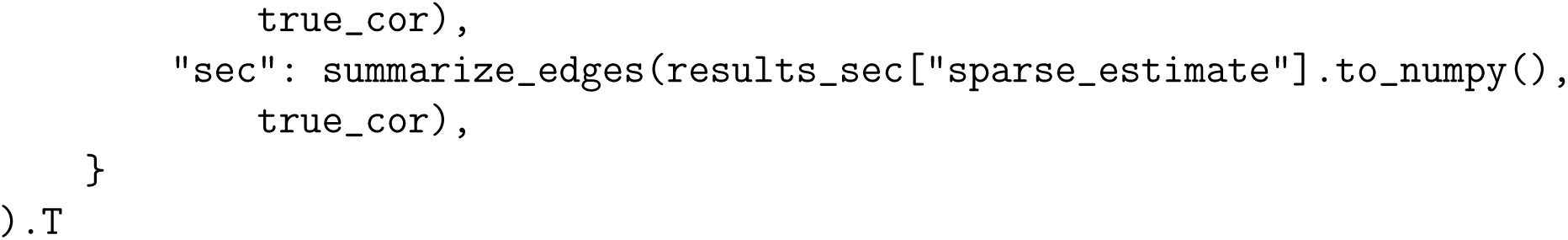

**CRITICAL:** The feature matrix and metadata must refer to the same samples. Misaligned sample identifiers will lead to incorrect covariate adjustment and unreliable correlation estimates.

**Note:** The main outputs of the workflow are stored in user-defined Python objects. The trained model stores correlation estimates internally, and the notebook illustrates both `sparse_by_p(` and `sparse_by_sec()` for obtaining sparse correlation matrices.

**Note:** In this example, zeros are introduced by masking the lowest 30% of simulated values within each metabolite. This is intended only to mimic left-censoring or missingness often observed in metabolomics data^20^.

**Note:** The Quickstart keeps most MetVAE settings at their package defaults and changes only the arguments needed to point the model to the input tables and identify continuous and categorical covariates.

**Note:** This example uses `workers=1` for `get_corr()` and `sparse_by_sec()` because that set-ting is often more stable in notebook environments. Increase the number of workers if parallel computing is available and faster execution is needed.

**Note:** Because the example data are simulated, the notebook also compares the output of `sparse_by_p()` and `sparse_by_sec()` against the known true correlation structure. This comparison is useful for learning, but it is not available for real experimental datasets.

## Key resources table

| REAGENT or RESOURCE | SOURCE | IDENTIFIER |
| --- | --- | --- |
| **Software and algorithms** |  |  |
| Python | Python Software Foundation | https://www.python.org/ |
| Jupyter Notebook | Project Jupyter | https://jupyter.org/ |
| NumPy | NumPy official website or conda-forge | https://numpy.org/ |
| pandas | pandas official website or conda-forge | https://pandas.pydata.org/ |
| seaborn | seaborn official website or conda-forge | https://seaborn.pydata.org/ |
| SciPy | SciPy official website or conda-forge | https://scipy.org/ |
| PyTorch | PyTorch official website or conda-forge | https://pytorch.org/ |
| NetworkX | NetworkX official website or conda-forge | https://networkx.org/ |
| Matplotlib | Matplotlib official website or conda-forge | https://matplotlib.org/ |
| MetVAE | PyPI | https://pypi.org/project/metvae/ |
| Cytoscape | Cytoscape project | https://cytoscape.org |
| Java | Java official website | https://www.java.com/ |
| **Other** |  |  |
| Apple M2 Max with 64 GB RAM | N/A | N/A |
| Enhanced Graphics Cytoscape App | Available from Apps > App Manager inside Cytoscape | N/A |

## Materials and equipment setup

### Install and load software package dependencies

Install Python version *≥* 3.9 on your computer. Then install and load the required Python dependencies for data import, data format manipulation, running MetVAE, and visualization. To generate molecular network (MN) visualizations, install Java 11 or later and download Cytoscape v3.9.1 or later. This protocol can be run on most laptop computers (the hardware used is listed in the key resources table).

### Step-by-step method details

The following worked example uses an untargeted metabolomics dataset from a hepatocellular carcinoma (HCC) mouse study to illustrate the full computational workflow. Specifically, the dataset derives from a study by Shalapour et al.^21^ in which MUP-uPA mice, a transgenic model predisposed to liver pathology, were fed a high-fat diet (HFD) for 3, 6, or 11 months to induce metabolic dysfunction-associated steatotic liver disease (MASLD), non-alcoholic steatohepatitis (NASH), and HCC, respectively.

In that study, HFD was identified as a major risk factor for HCC development and was associated with microbial dysbiosis and altered fecal metabolites. This biological setting provides a useful demonstration case for MetVAE because it contains the types of coordinated metabolite variation that correlation-based networking is designed to recover. In the steps below, we use this HCC dataset to show how to import and harmonize the data, fit MetVAE, estimate sparse metabolite correlations, export a GraphML network, and generate the final Cytoscape visualization.

### Import and harmonize metabolomics data for the HCC study

This major step imports the MS1 abundance table, (optional) MS2 annotation table, and sample metadata, then harmonizes sample identifiers and excludes samples not used in the downstream MetVAE analysis.

**Timing:** *<* **1 min**

1. Import the required Python libraries.

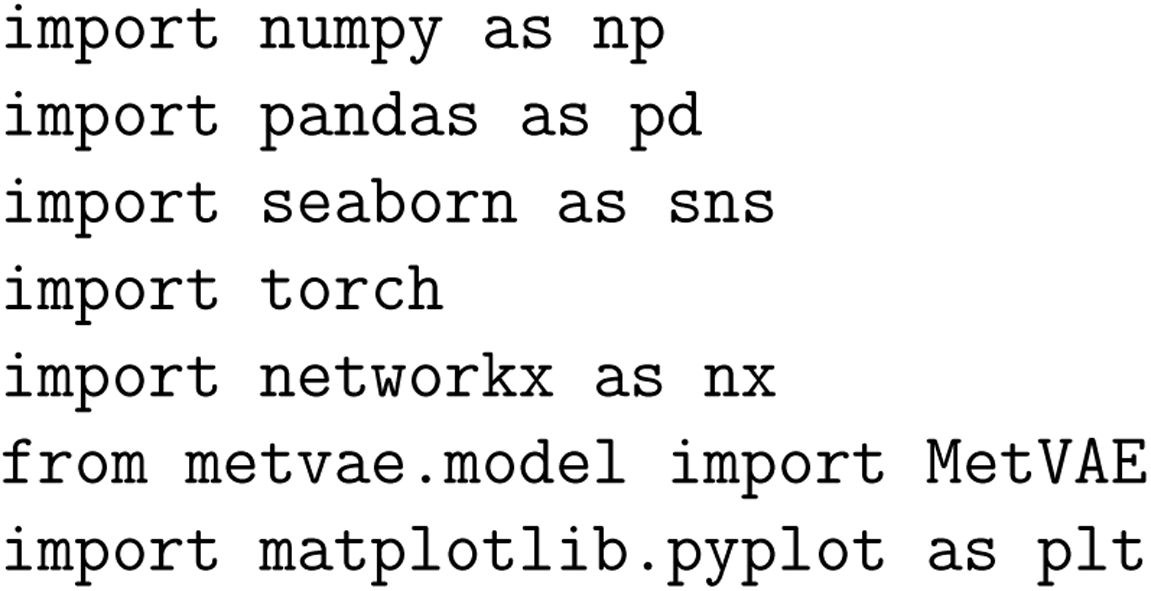
2. Import the MS1 feature table, MS2 table, and sample metadata.

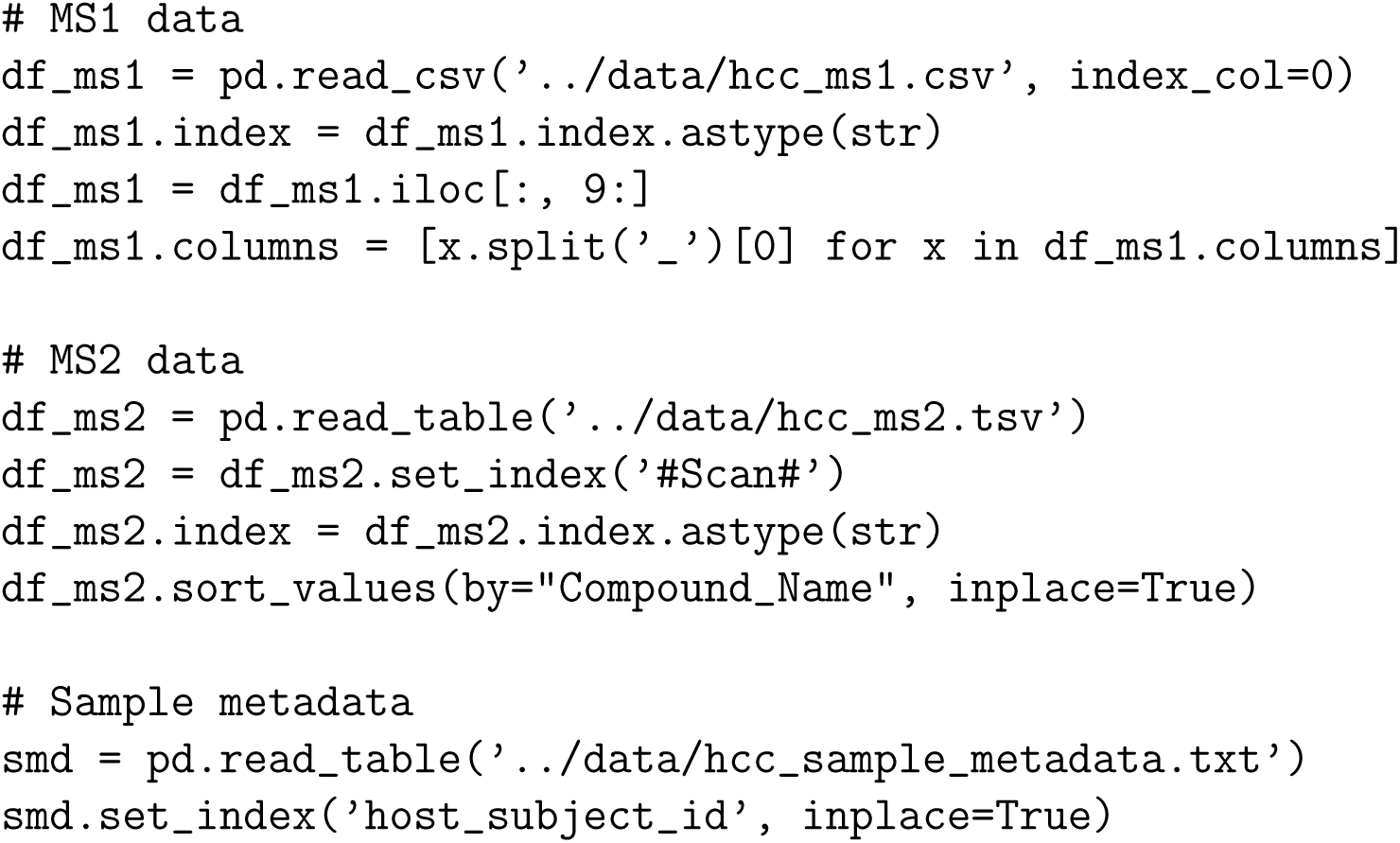
3. Standardize metadata fields used later in the analysis.

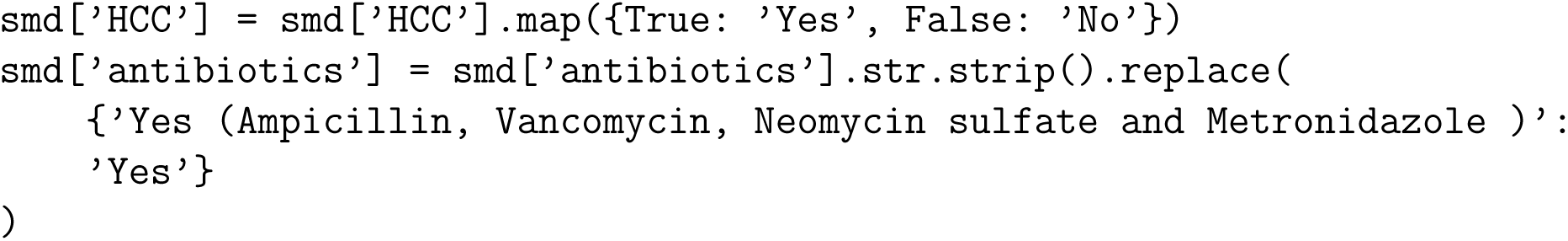
4. Remove duplicated MS1 sample columns, exclude samples exposed to antibiotics, and retain only sample identifiers shared between the MS1 table and metadata.

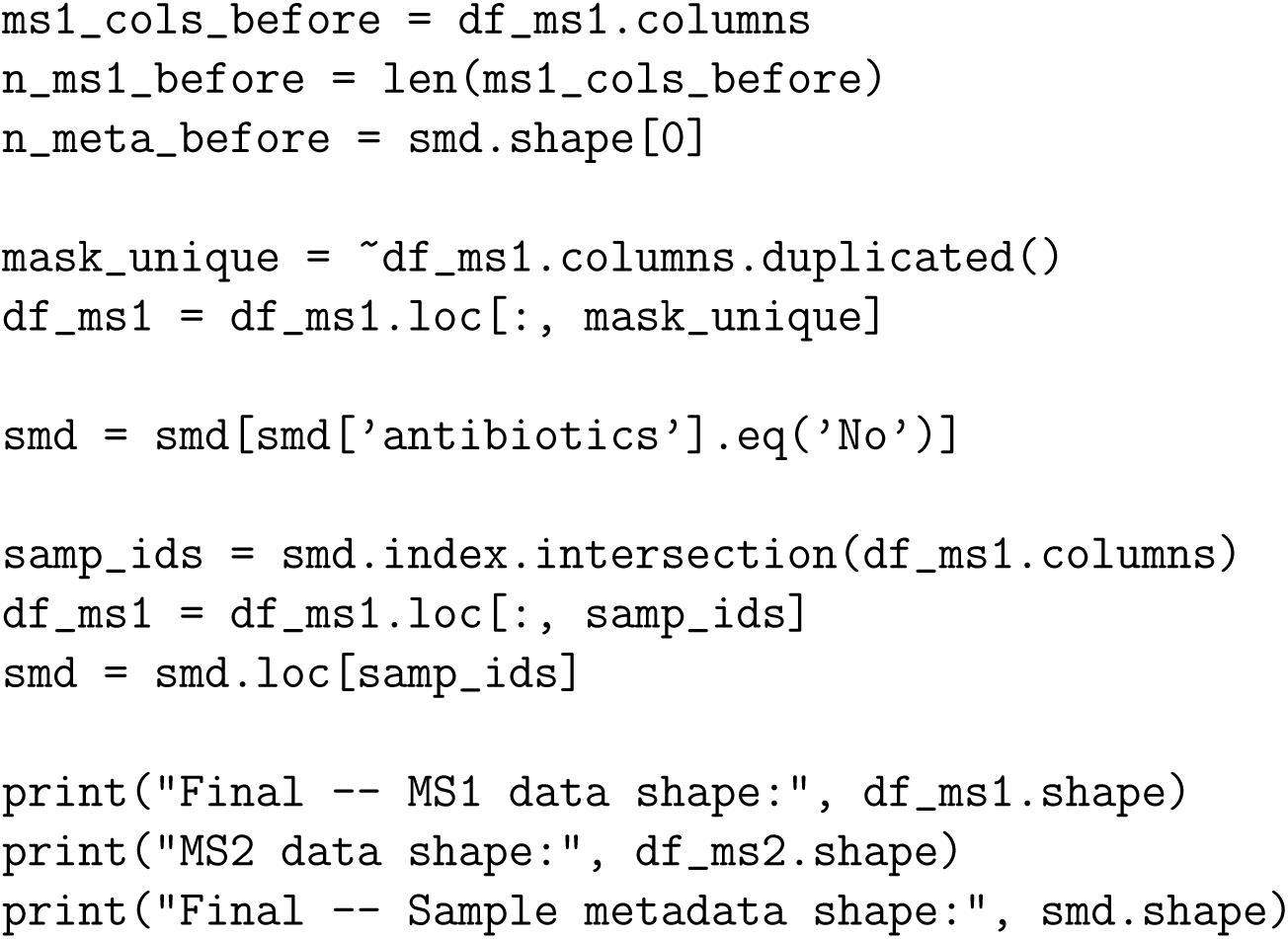

**CRITICAL:** Align the sample identifiers in the feature table and metadata before fitting MetVAE. Sample mismatches will propagate into incorrect covariate adjustment and downstream network construction.

**Note:** In this example, samples with `antibiotics != ’No’` were excluded before fitting the model. If a different study design is used, replace this filtering rule with the selection criteria appropriate for that dataset.

**Note:** The example output reported 411 aligned samples after removing one duplicated MS1 column, excluding 27 metadata entries with antibiotic exposure, and dropping non-overlapping sample identifiers.

### Construct the abundance matrix and initialize the MetVAE model

This major step converts the MS1 table into the sample-by-feature matrix required by MetVAE, filters sparse features, and initializes the MetVAE model used for correlation estimation.

**Timing: 2-5 min, depending on the computer’s hardware**

5. Transpose the MS1 table so that rows correspond to samples and columns correspond to metabolite features, then remove highly sparse features.

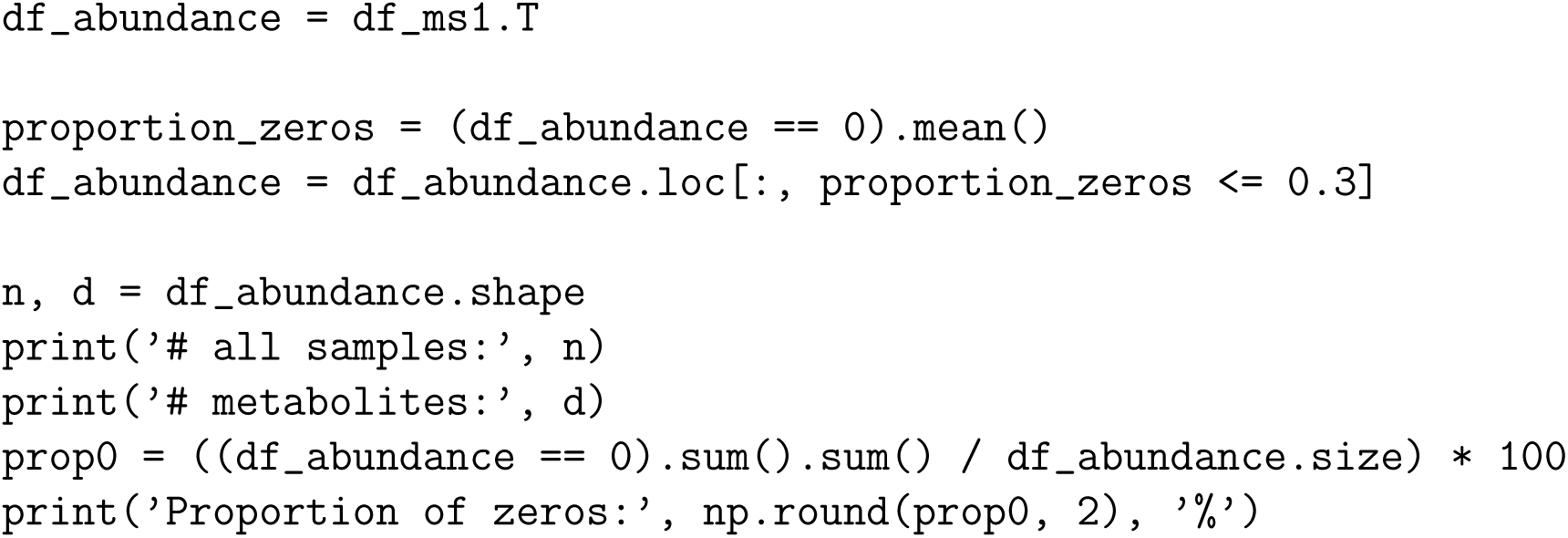
6. Initialize the MetVAE model.

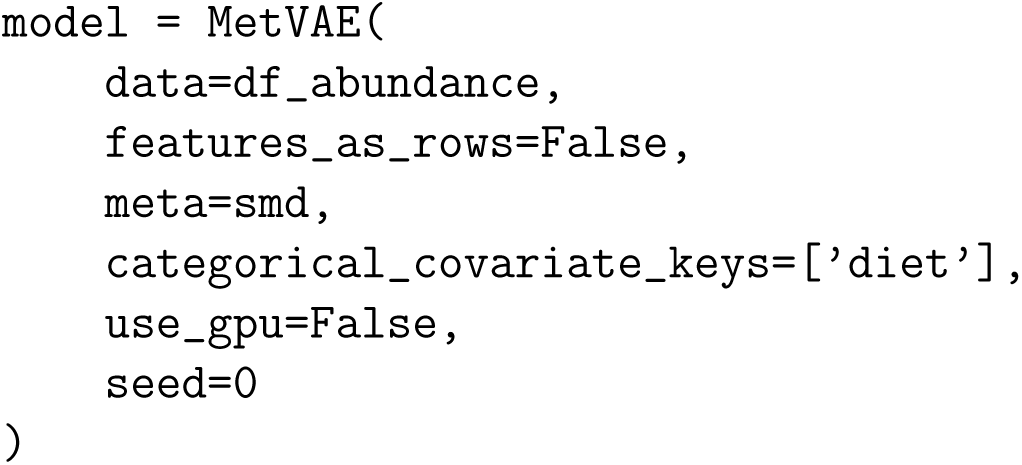
7. Train the model from scratch or load a previously saved checkpoint.

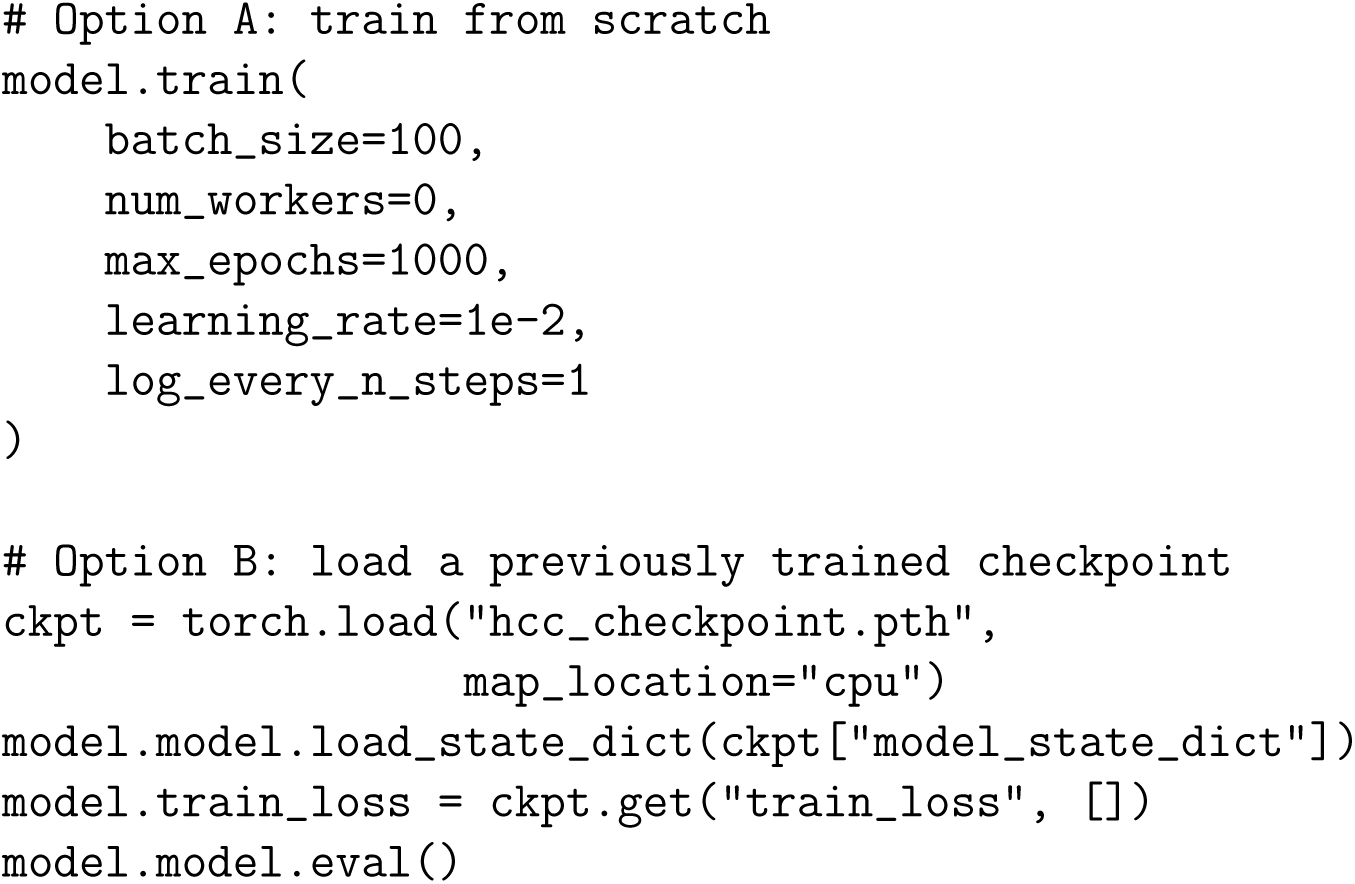

**CRITICAL:** When `features as rows=False`, MetVAE expects the abundance matrix to have samples in rows. If the matrix orientation is reversed, either transpose the table or change this argument accordingly.

**Note:** `categorical covariate keys=[’diet’]` instructs MetVAE to model diet as a categorical covariate during training. Add additional covariates here if they should be adjusted in the latent model.

**Note:** Users can specify `feature zero threshold` in `MetVAE` to remove features whose zero proportion exceeds a user-specified threshold during preprocessing, and `sample zero threshold` optionally removes samples with excessive zeros. In the HCC example, feature filtering was performed manually before model initialization by retaining features with *≤* 30% zeros.

**Note:** In the `MetVAE` function, users can specify `latent dim`, the dimension of the latent representation learned by the VAE. By default, it is set to the smaller of the sample size and the number of features. We used the default in this example and therefore did not specify this parameter.

**Note:** `use gpu=False` forces computation on CPU. Set `use gpu=True` only when a compatible CUDA-enabled GPU is available.

**Note:** `batch size`, `max epochs`, `learning rate`, and `log every n steps` control mini-batch training, total training length, optimizer step size, and logging frequency, respectively. The optimization process is summarized in Figure 2, which shows both the raw training loss and a moving-average trend across epochs.

**Figure 2.**
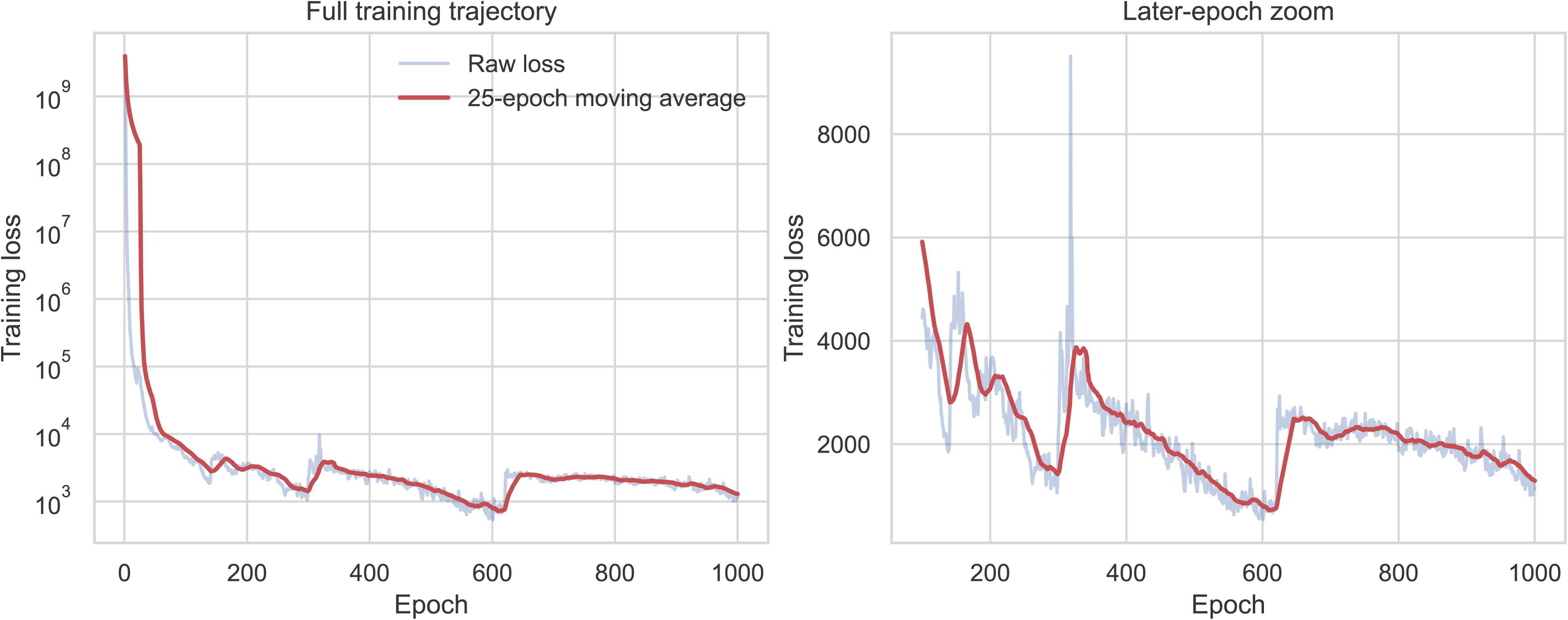
Training-loss profile for the MetVAE model fitted to the HCC dataset. Training-loss trajectories during MetVAE optimization for the HCC example. (Left panel) Full training trajectory across 1,000 epochs. The light-blue curve shows the raw training loss, and the red curve shows the 25-epoch moving average. The y axis is plotted on a logarithmic scale to highlight the rapid early loss decrease followed by slower stabilization. (Right panel) Zoomed view of later epochs on a linear scale, showing the same raw-loss and moving-average curves in greater detail. This panel helps assess convergence behavior after the initial training phase and illustrates the residual fluctuations around the overall downward trend.

### Estimate the sparse correlation matrix

This major step generates a correlation estimate from multiple imputations of censored zeros and then applies sparsification to retain only interpretable edges. MetVAE provides two sparsi-fication strategies, `sparse by p()` and `sparse by sec()`. In this HCC example, we proceed with `sparse by p()` for downstream analysis.

**Timing: 4-6 min, depending on the computer’s hardware**

8. Estimate the correlation matrix from repeated imputations.

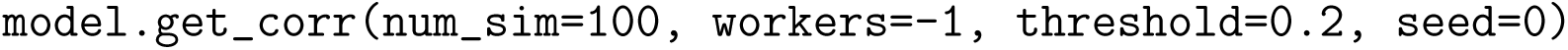

**CRITICAL:** Run `model.get corr()` before calling `sparse by p()` or `sparse by sec()`. The sparsification step relies on the correlation estimate stored inside the trained model object.

**Note:** `num sim` controls the number of multiple imputations used for correlation estimation. Larger values may improve stability at the cost of additional computation time. The default is 100.

**Note:** The `threshold` argument in `get corr()` is used as an internal hard correlation threshold used during correlation computation. That is, correlations less than `threshold` will be considered 0 in the current iteration. The default is 0.2.

**Note:** `workers` controls the number of CPU workers used during correlation estimation and is ignored when GPU-based imputation is used. The default value is -1, which will use all available CPUs.

9. Option A: apply *P* -value-based sparsification with `sparse by p()`.

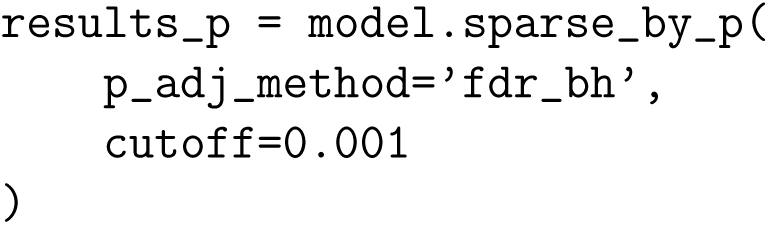

**Note:** `sparse by p()` is the more direct of the two sparsification options. It starts from the estimated correlation matrix, asks which metabolite pairs have enough evidence to keep, adjusts for multiple testing, and sets the remaining weakly supported edges to zero. In this example, we use Benjamini-Hochberg^22^ adjustment with `p adj method=’fdr bh’` and retain only edges with adjusted *P* -values below 0.001. Because many pairwise tests are evaluated from the same dataset, the adjusted *P* -values should be interpreted mainly as an exploratory filtering tool.

**Note:** In this protocol, we continue with `sparse by p()` for downstream analysis because we want to maximize power and generate hypotheses from a large untargeted dataset. SEC remains a useful alternative when a sparser and positive-definite estimate is preferred.

10. Option B: apply SEC-based sparsification with `sparse by sec()`.

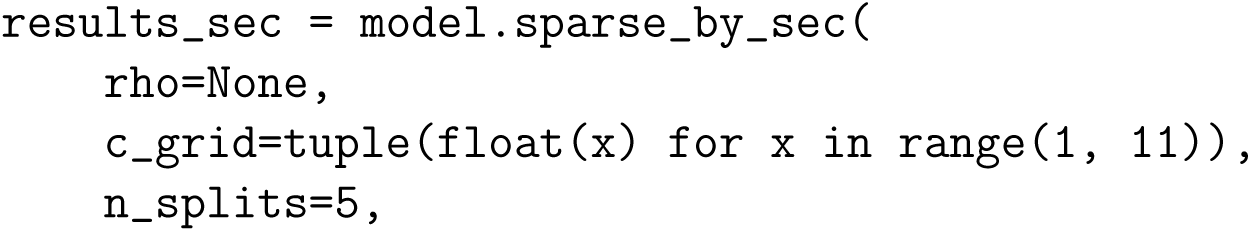
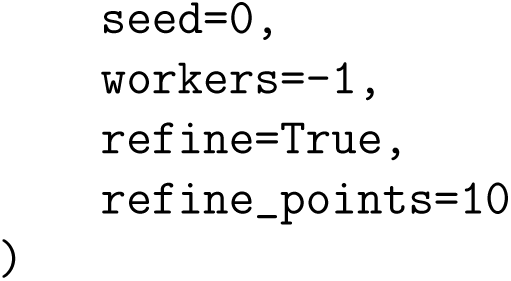

**Note:** `sparse_by_sec()` uses the Sparse Estimation of Correlation (SEC) algorithm^23^ to produce a sparse positive-definite correlation matrix. Conceptually, SEC does not test edges one by one. Instead, it estimates the whole correlation matrix while shrinking small or unstable correlations toward zero, which often gives a cleaner and more conservative network. This option is especially useful when users want a sparse matrix that is numerically well behaved for downstream analysis.

**Note:** The main tuning parameter in SEC is `rho`, which controls how strongly small correlations are shrunk toward zero. Larger values generally produce sparser networks. If a fixed `rho` is supplied, SEC runs once at that penalty value. If `rho=None`, as in the example above, MetVAE selects the penalty automatically by cross-validation.

**Note:** When SEC is run with automatic penalty selection, `c_grid` defines the coarse set of candidate scaling constants used to generate penalties according to 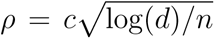, where `c` come from `c_grid`, *d* is the number of metabolite features, and *`n`* is the number of samples. The argument `n_splits` sets the number of cross-validation folds used to compare these candidate penalties; larger values use more repeated train-validation splits but increase computation time. The argument `seed` makes fold assignment reproducible, and `workers` controls how many CPU workers are used to evaluate candidates. The default value is -1, which will use all available CPUs. If `refine=True`, MetVAE performs a second, narrower search around the best coarse penalty found in the first pass. The argument `refine_points` determines how many evenly spaced candidate values are evaluated in this zoomed-in search, with larger values giving a finer local search at the cost of additional computation.

**Note:** The remaining SEC arguments mainly control optimization and post-processing. In particular, `tol` and `max_iter` determine when optimization stops, `restart` and `line_search_ap` help stabilize the accelerated optimization routine, `epsilon` is used during positive-semidefinit calibration, and `threshold` sets a final hard cutoff so that very small SEC estimates are set to zero. The arguments `delta`, `n_samples`, and `c_delta` are used when the internal small-correlation threshold is chosen automatically. For most users, the default settings are a reasonable starting point.

11. For the HCC example, proceed with the `sparse_by_p()` output for downstream analysis.

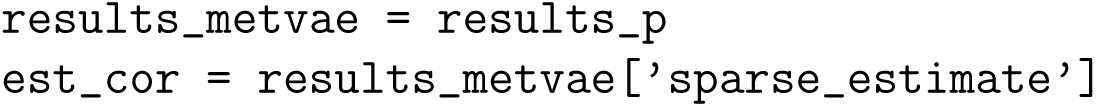

### Export a correlation network as GraphML

This major step thresholds the MetVAE edge list, transfers node attributes from the original molecular network, and writes a GraphML file for downstream visualization.

**Timing: 1-2 min**

12. MetVAE can export GraphML directly from the sparse correlation matrix. We keep that code here for reference, but we do not run it because the analysis below adds correlation edges onto an existing cosine-similarity graph.

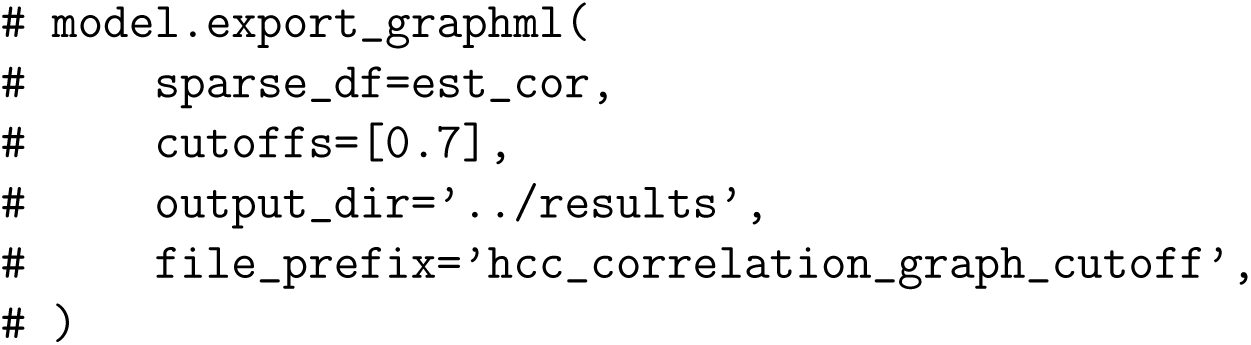
13. Load the cosine-similary based molecular network and the MetVAE edge list.

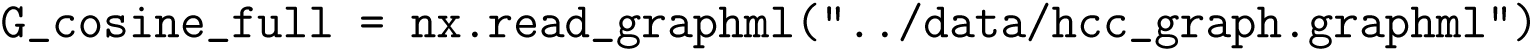
14. Define helper functions for converting the sparse correlation matrix into a graph-ready edge table and GraphML-ready network.

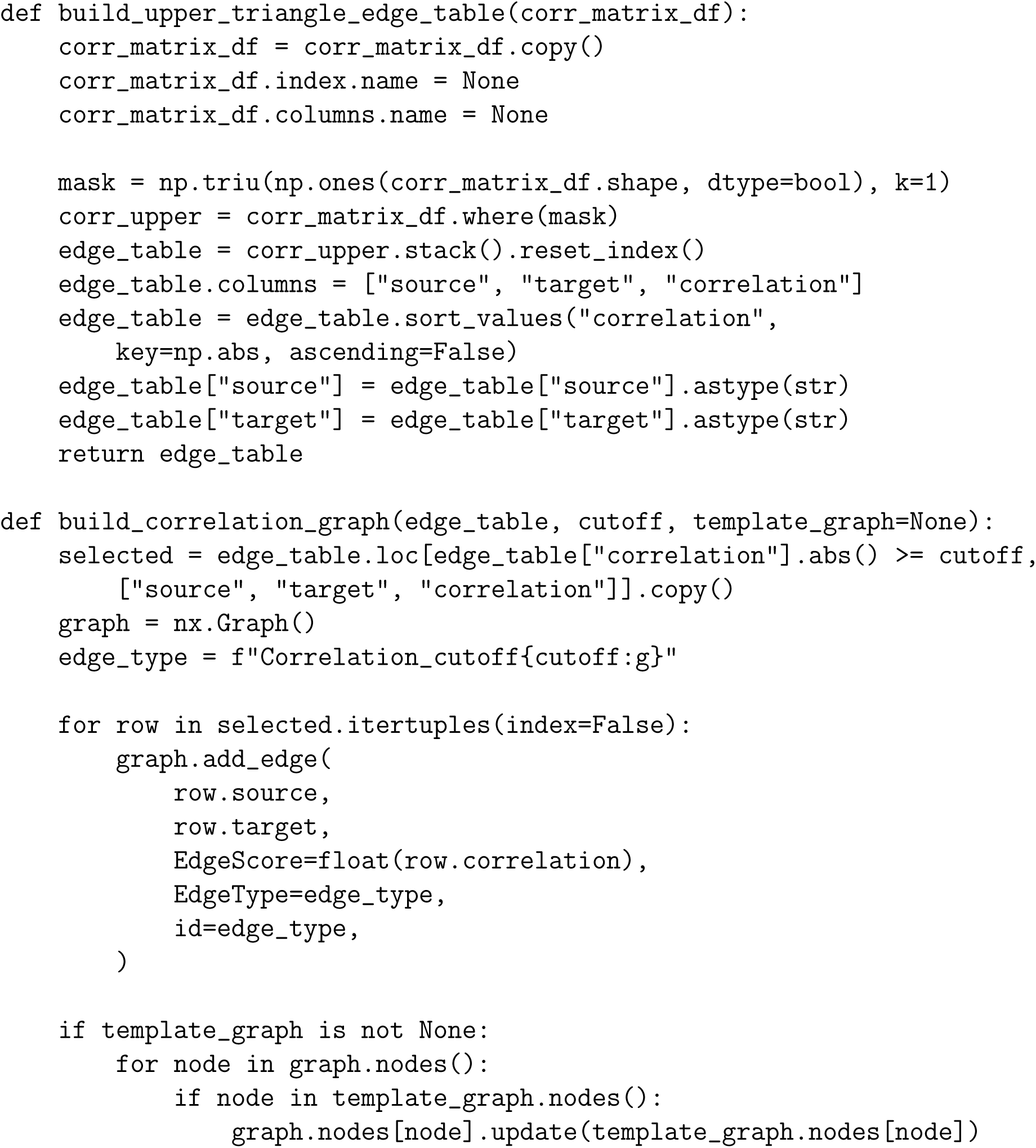
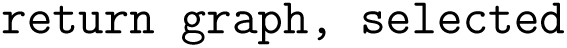

**Note:** The helper function `build_upper_triangle_edge_table()` converts the square metabol metabolite correlation matrix into a simple edge table. Because the correlation matrix is symmetric, the function keeps only the upper triangle so that each metabolite pair appears once rather than twice. It then stores the result as a long table with `source`, `target`, and `correlation` columns and sorts the rows by absolute correlation strength.

**Note:** The helper function `build_correlation_graph()` then converts that edge table into a NetworkX graph that is ready for GraphML export. It applies a user-defined correlation cutoff, keeps only metabolite pairs above that threshold, adds them as graph edges, and stores edge attributes such as correlation strength and edge type. If a template graph is provided, the function also copies over node attributes from that graph. This step is useful when the final figure should display structural annotations, feature names, or sample-level summaries already curated elsewhere. In this example, the template graph is the cosine-similarity molecular network, which allows the MetVAE correlation graph to inherit curated node annotations for downstream visualization.

15. Run the helper functions to build the MetVAE correlation graph.

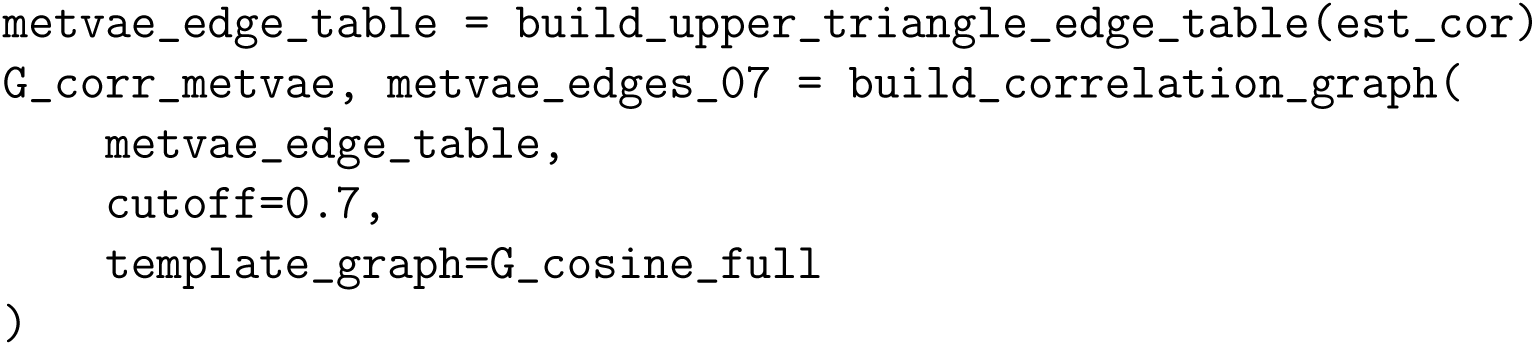

**Note:** In this example, a cutoff of `|correlation| >= 0.7` was used to define the exported network. Adjust this threshold to balance network density and interpretability for the dataset being analyzed.

16. Export the MetVAE correlation graph

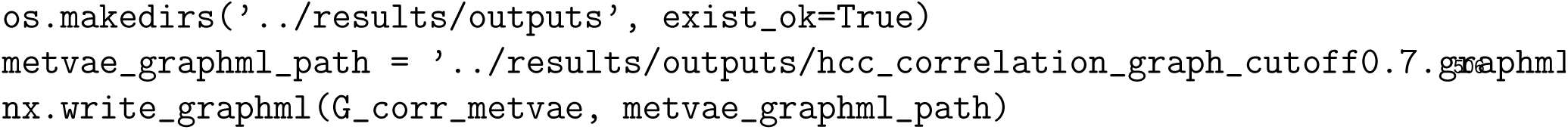

**Note:** The exported GraphML file is intended for downstream visualization in Cytoscape or other network-analysis software.

### Molecular network visualization in cytoscape: import, styling, color mapping, and export

The molecular network (MN) was visualized using Cytoscape (v3.9.1 or later). The GraphML network exported in the previous step was imported into Cytoscape, styled using the built-in style editor, augmented with group-level pie charts using the Enhanced Graphics app, and saved as a Cytoscape session file (`.cys`). The steps below describe a reproducible workflow for import, layout, styling, color mapping, chart overlay, and image export.

**Timing: 30-45 min**

17. Before opening Cytoscape, confirm that the GraphML file contains the node and edge attributes needed for visualization.

**CRITICAL:** The GraphML file should include a node attribute for the group label (for example, compound class or lipid family), an edge attribute for the correlation sign or direction, and numeric node attributes for covariates (in our example, normal chow (NC) and high-fat diet (HFD) abundance) if pie charts will be added later. If these attributes are missing, the styling steps below will not work as written.

18. Install the Enhanced Graphics app if it is not already available. Open Cytoscape, go to *Apps > App Manager*, search for *Enhanced Graphics*, install it, and restart Cytoscape.
19. Launch Cytoscape and import the GraphML file by going to *File > Import > Network from File*. Select the exported `.graphml` file and click *Open*.

**Note:** Cytoscape reads `.graphml` files natively, so no additional import plugin is required. Make sure the file extension is `.graphml`; files saved with other extensions such as `.xml` or `.gml` may not import correctly through this menu.

20. Confirm that the imported network appears correctly by checking the node and edge counts and inspecting the Node Table and Edge Table to verify that the expected attributes are present.

**Note:** If the network canvas appears blank after import, inspect the GraphML file in a text editor and check for formatting or encoding problems before re-importing.

21. Apply a layout by going to *Layout > Perfuse Force Directed Layout*. This provides a readable starting point and can be adjusted later without changing any styling choices.
22. Open the *Style* panel and choose the Cytoscape style named *Sample 1* as a clean starting point.

**CRITICAL:** If *Sample 1* is not available, restore or import the default Cytoscape styles before continuing.

23. Color nodes by group. In the *Node* tab of the *Style* panel, expand *Fill Color*, select the node attribute containing group labels, and use *Discrete Mapping* to assign a distinct fill color to each group.
24. Color edges by correlation direction. In the *Edge* tab of the *Style* panel, expand *Stroke Color (Unselected)*, choose the edge attribute encoding correlation sign or direction, and use *Discrete Mapping* to assign colors such as pink (#FF69B4) for positive correlations and blue (#4169E1 or #1E90FF). for negative correlations.

**CRITICAL:** Check the exact values stored in the edge attribute column before mapping colors. The mapping keys in Cytoscape must match those values exactly.

25. Add black borders to all nodes by setting *Border Paint* to black as a default value and adjusting *Border Width* to approximately 1.5–2.5, depending on network density.
26. Set node labels by mapping the *Label* property to the attribute containing compound or feature names using *Passthrough Mapping*. Then adjust label font size and label color so that names remain legible without overwhelming the graph.

**Note:** If labels are too long for practical display, create a shortened label column in the GraphML file before import and map labels to that field instead.

27. Adjust node size and edge width to improve readability. For most networks, node sizes in the range of 35–60 pixels and edge widths in the range of 1.0–3.0 pixels provide a good starting point.
28. Add group-level pie charts to each node using the Enhanced Graphics app. First, verify that the Node Table contains numeric abundance columns for NC and HFD. Then add a new string column (for example, `pie chart`) and populate it with the Enhanced Graphics syntax specifying the NC and HFD columns and their corresponding colors.

**CRITICAL:** If the NC and HFD abundance columns are missing from the Node Table, go back and regenerate the GraphML file with those attributes included before attempting this step.

29. Map the new pie-chart column to *Image/Chart 1* in the *Node* style properties using *Passthroug Mapping*. Confirm that each node now displays a pie chart with green representing NC and red representing HFD. The final Cytoscape-rendered network should resemble the example shown in Figure 3.

**Figure 3.**
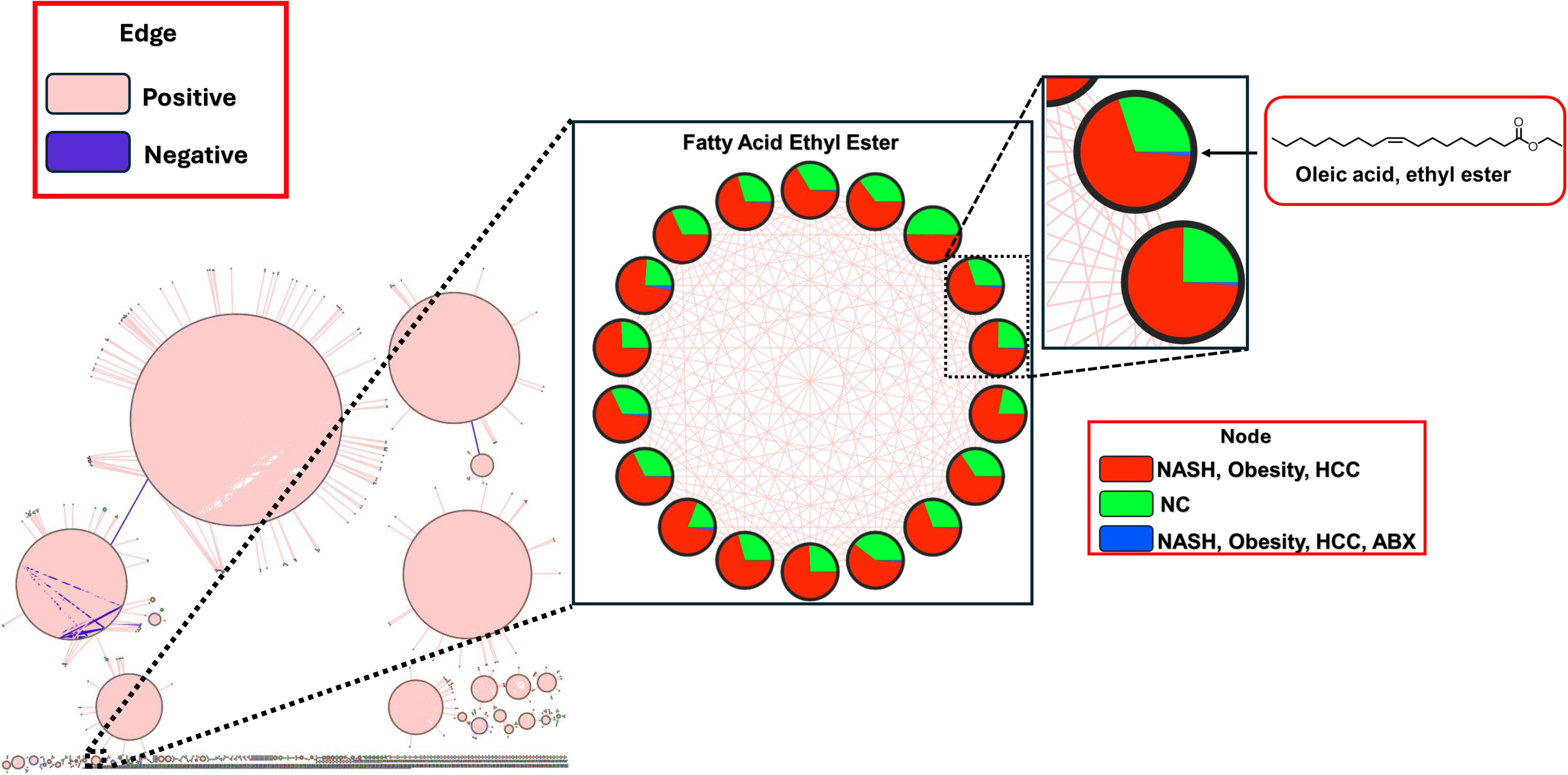
Correlation-based molecular network of HCC samples (circular layout). Edges represent pairwise correlations with absolute value *≥* 0.7 (i.e., both positive and negative correlations are included). Each node is shown as a pie chart, where the colors indicate the average metabolite abundance across samples within each group. The phosphocholine (PC) lipid cluster is highlighted; for these nodes, the pie charts specifically display the average metabolite abundance in the normal chow (NC) and high-fat diet (HFD) groups.

**Note:** The same chart-specification string should be present in each row of the new chart column, while Cytoscape uses the node-specific abundance values to determine slice sizes.

30. Save the Cytoscape session as a `.cys` file using *File > Save Session As*.

**CRITICAL:** Save the final styled network as a `.cys` session file. This is the only format that preserves the full network, layout, visual styles, color mappings, pie-chart overlays, and labels.

31. Reopen the saved `.cys` file to confirm that all styles and chart overlays are retained. If the session reloads correctly, export the final network to SVG or high-resolution PNG for figure preparation.

**Note:** The exact color palette, label density, and final layout can be adjusted to match journal figure requirements, but the general workflow above preserves the main visual logic of the HCC example.

### Expected outcomes

Applying this protocol to the HCC metabolomics dataset is expected to produce a filtered sample-by-feature abundance matrix, a trained MetVAE model, a sparse metabolite-metabolite correlation estimate, and an exported GraphML network that can be visualized in Cytoscape. For the HCC example presented here, preprocessing yields 411 aligned samples and 7,217 metabolite features after sample matching and feature filtering. After correlation estimation and downstream thresholding at *|r| ≥* 0.7, the exported network contains a reduced set of strongly associated edges that can be overlaid with node attributes from the original molecular network.

In this HCC use case, the correlation network highlighted coordinated elevation of multiple lipid classes in HFD animals. Beyond expected increases in lipids such as phosphatidylcholines, MetVAE identified a co-varying module containing monoacylglycerols (MAGs), free fatty acids, bile acids, and fatty acid ethyl esters (FAEEs). This pattern was not apparent in conventional cosine-similarity molecular networking (cosine cutoff 0.7) because these metabolite classes are structurally distinct (Figure 4). The observed correlation pattern suggests a putative functional linkage consistent with known steps in gut–liver metabolism: HFD-associated dysbiosis may promote malabsorption of phospholipids, MAGs, and free fatty acids into the colon, where bacterial degradation and fermentation can generate ethanol that subsequently esterifies with fatty acids to form FAEEs. Notably, FAEEs are the same toxic metabolites implicated in alcohol-associated liver disease^24^, despite the absence of alcohol intake in this model. Together, these associations point to a previously unrecognized possibility for an endogenous ”auto-brewery” route in which dietary fat, mediated by microbiome alterations, contributes to alcohol-associated hepatotoxic chemistry. This hypothesis may also help contextualize the long-noted overlap between MASLD and alcohol-associated liver disease, which can exhibit highly similar histological features, including steatosis, steatohepatitis, hepatocellular ballooning, Mallory-Denk bodies, and progression to cirrhosis and HCC, despite ostensibly distinct etiologies^25–27^.

**Figure 4.**
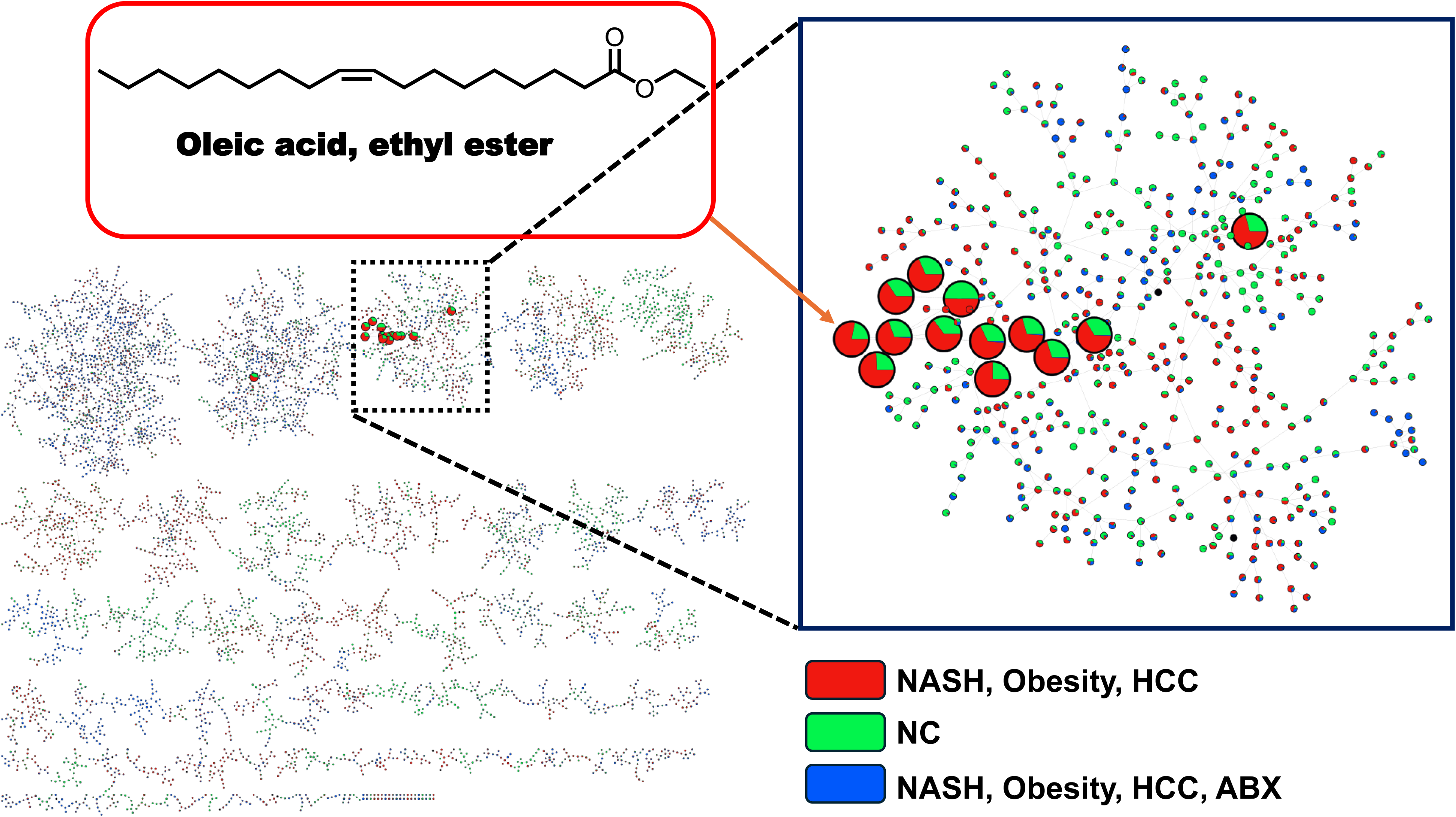
Structural similarity (cosine-based) molecular community network with highlighted nodes corresponding to the cluster shown in Figure 3. Each node is displayed as a pie chart, with colors representing the average metabolite abundance across samples in the NASH-obesity-HCC, normal chow (NC), and NASH-obesity-HCC with antibiotics groups. Fatty acid ethyl esters (FAEEs) group within a single molecular community based on structural similarity. However, their coordinated association pattern is not clearly apparent in this network, as the FAEE nodes are embedded within a large cluster that includes a diverse range of fatty acid–related structures.

This example illustrates how correlation-based networking can reveal putative functional relationships across molecular classes and generate biologically testable hypotheses. More generally, the final network should reveal clusters of co-varying metabolite features suitable for hypothesis generation, prioritization of candidate modules, and comparison with cosine-similarity-based molecular networking. Depending on the dataset and sparsification strategy, the exact number of retained edges and modules will vary.

After Cytoscape styling, the final network should also provide a visually interpretable summary of the data, including nodes colored by metabolite group, black node borders, positive and negative edges shown with distinct colors, readable node labels, and group-level pie charts showing relative NC and HFD abundance within each node. The saved `.cys` session should re-open with these settings intact, allowing the figure to be reproduced and refined later if needed.

## Quantification and statistical analysis

MetVAE is designed to address four common statistical challenges in untargeted metabolomics data. First, compositionality is handled by transforming the data onto a centered log-ratio (CLR) scale, which makes relative-abundance measurements more suitable for correlation analysis. Second, missingness is addressed through VAE-based reconstruction with multiple imputations of censored or zero-valued entries before correlation estimation. Third, confounding is handled by incorporating observed covariates directly into the model so that downstream correlations are less driven by technical or biological nuisance structure. Fourth, high dimensionality is addressed by combining latent-variable modeling with downstream sparsification, which enables estimation of interpretable metabolite-metabolite relationships in datasets with many more features than would be tractable using naive pairwise analysis alone. Interested readers should consult the Supplementary Information for full methodological details and model derivations.

Sparse correlation matrices can be obtained in MetVAE using either `sparse_by_p()` or `sparse_b` The `sparse_by_p()` option applies Fisher’s *z*-test to the estimated correlations, adjusts the resulting *P* -values for multiple testing, and sets non-significant entries to zero. In this protocol, we used Benjamini-Hochberg^22^ adjustment with `p_adj_method=’fdr_bh’` and `cutoff=0.001`. The `sparse_by_sec()` option instead uses Sparse Estimation of Correlation (SEC)^23^ algorithm to obtain a sparse positive-definite correlation matrix, either with a user-specified penalty parameter or with cross-validated penalty selection. Both approaches produce sparse correlation estimates, but for the HCC example we used `sparse_by_p()` for downstream analysis because we wanted to maximize power for hypothesis generation.

For covariate adjustment in the HCC analysis, diet was incorporated directly into the MetVAE model as a categorical covariate distinguishing high-fat diet (HFD) and normal chow (NC). Specifically, we adjusted for diet effects on the CLR-transformed metabolite abundances by including diet as a covariate and removing the estimated NC effect. This procedure yields diet-adjusted metabolite levels centered at the HFD mean for all mice, which allows downstream correlation analysis to focus on residual co-variation after accounting for the modeled diet effect.

For downstream network export, the sparse correlation estimate was converted to an edge list and then filtered using an edge-retention rule of *|r| ≥* 0.7. This additional thresholding step was used to generate a network of strongly associated metabolite pairs for GraphML export and Cytoscape visualization. Summary statistics were recorded throughout preprocessing and network construction to document the analysis. In the HCC example, these included the number of aligned samples, the number of retained metabolite features after filtering, the overall proportion of zeros in the abundance matrix, and the number of retained edges after applying the final correlation cutoff. These summaries provide a compact check on dataset size, sparsity, and network density before biological interpretation.

## Limitations

This protocol is designed for cross-sectional metabolomics data and does not explicitly model longitudinal or repeated-measures structure. When multiple samples are collected from the same subject over time, within-subject dependence and temporal dynamics may influence correlation estimates in ways that are not captured by the current workflow. In such settings, additional modeling would be needed to account for subject-specific effects and time-dependent covariance structure.

As with any correlation-based analysis, the inferred network describes statistical association rather than causal relationships. Strongly correlated metabolites may arise from direct biochemical coupling, shared upstream regulation, common environmental exposure, batch-related structure, or other latent factors. Therefore, network edges should be interpreted as candidates for follow-up rather than as evidence of direct mechanistic interaction.

The protocol also depends on the quality of upstream metabolomics preprocessing. Errors in peak detection, alignment, feature integration, annotation transfer, or sample metadata harmonization can propagate into the abundance matrix and affect downstream correlations. In addition, although MetVAE is designed to address compositionality, missingness, confounding, and high dimensionality, the final network can still be sensitive to user-defined choices such as feature filtering thresholds, covariate specification, sparsification strategy, and correlation cutoffs used for GraphML export.

Finally, this workflow is most useful for hypothesis generation and network prioritization rather than definitive biological inference. Sparse correlation modules can highlight coordinated metabolite behavior and reveal patterns that complement conventional molecular networking, but biological interpretation still depends on metabolite annotation quality, external validation, and follow-up experimental or orthogonal computational analyses.

## Troubleshooting

### Problem 1: Feature table and metadata do not align (Step 4)

After filtering, the number of samples in the abundance table and metadata may not match, or the sample order may differ between the two objects.

### Potential solution

Confirm that the sample identifiers use the same format in both files before subsetting. Remove duplicated sample columns from the MS1 table, standardize identifier formatting, and retain only the intersection of sample identifiers shared between the abundance table and metadata. After alignment, verify that the abundance matrix columns and metadata rows refer to the same samples in the same order before initializing MetVAE.

### Problem 2: Too many features are removed during preprocessing (Step 5)

After applying feature filtering, the retained feature set may be unexpectedly small, reducing network coverage and limiting downstream interpretation.

### Potential solution

Inspect the zero proportion across features before filtering and evaluate whether the threshold is too stringent for the dataset. In the HCC example, features with more than 30% zeros were removed manually before model fitting. If your dataset is sparser, consider testing a less restrictive threshold while monitoring whether the retained feature set still yields stable training and interpretable correlation structure.

### Problem 3: Covariate adjustment appears to remove biological signal of interest (Step 6)

Including covariates in the model may attenuate correlation patterns that appear biologically relevant, especially when the covariate is closely related to the phenotype being studied.

### Potential solution

Reconsider whether the variable should be treated as a nuisance covariate, a variable of interest, or a stratification factor. In the HCC example, diet was included deliberately to generate diet-adjusted metabolite levels centered at the HFD mean. For other studies, it may be more appropriate to omit a variable from adjustment, analyze groups separately, or compare adjusted and unadjusted results to determine how strongly that variable drives the network structure.

### Problem 4: Model training is unstable or does not converge well (Step 7)

Training loss may decrease slowly, fluctuate strongly, or plateau at a high value, making the learned representation difficult to interpret.

### Potential solution

Check that the input matrix orientation is correct and that the data are being passed with samples in rows when `features_as_rows=False`. Review the training settings, especially `learning_rate`, `batch_size`, `max_epochs`, and `latent_dim`. If needed, reduce the learning rate, increase the number of epochs, or test a smaller latent dimension. Plotting the training loss, as shown in Figure 2, can help determine whether the model is still improving or has become unstable.

### Problem 5: The sparse correlation matrix is too dense or too sparse (Steps 8–11)

After sparsification, the resulting network may contain too many edges to interpret clearly or too few edges to reveal meaningful modules.

### Potential solution

Adjust the sparsification settings according to the analysis goal. For `sparse_by_p()`, modify the adjusted *P* -value cutoff to be more or less stringent. For `sparse_by_sec()`, review the selected penalty or the candidate penalty grid used in cross-validation. In addition, the final edge-retention threshold used for network export, such as *|r| ≥* 0.7, can be relaxed or tightened to control final network density independently of the initial sparsification step.

### Problem 6: The exported GraphML file does not display correctly in Cytoscape (Steps 17–20)

The imported network may appear to have missing node attributes, unlabeled nodes, or unexpected edge styling in Cytoscape.

### Potential solution

Verify that node attributes were successfully copied from the original molecular-network GraphML file (if any) before export and that the final GraphML file was written without errors. Inspect a subset of nodes and edges in Python before export to confirm that key attributes such as feature identifiers, edge scores, and edge types are present. If Cytoscape still does not display the network as expected, re-import the GraphML file and confirm that the correct attribute columns are being used for node labels, color mapping, and edge styling.

### Problem 7: The expected Cytoscape style is not available (Steps 22–23)

The *Sample 1* style may be missing from the Cytoscape style list, making it difficult to reproduce the styling workflow described in this protocol.

### Potential solution

Restore or import the default Cytoscape styles before proceeding. If the Cytoscape installation was modified or the default styles were removed, reinstall them or choose another clean baseline style and then reapply the mappings described in this protocol.

### Problem 8: Node colors or edge colors do not update as expected (Steps 23–24)

Nodes or edges may remain gray or display the wrong colors after mapping is applied.

### Potential solution

Check that the correct attribute column has been selected and that the mapping type is appropriate. Use *Discrete Mapping* for group labels and correlation direction, and verify that the values stored in Cytoscape match the mapping keys exactly, including capitalization and spacing. If some items still fail to update, inspect the Node Table or Edge Table to identify unmatched values.

### Problem 9: Pie charts do not appear inside nodes (Steps 28–29)

The Enhanced Graphics overlay may fail to render, or nodes may appear without the expected NC/HFD pie charts.

### Potential solution

Confirm that the Enhanced Graphics app is installed and active, that the Node Table contains the NC and HFD abundance columns, and that the chart-specification column was created as a string field with valid syntax. Then map that column to *Image/Chart 1* using *Passthrough Mapping*. If charts still do not appear, inspect one node entry manually to confirm that the syntax string and referenced column names are correct.

### Problem 10: The Cytoscape session does not reopen with all visual settings preserved (Steps 30–31)

After reloading Cytoscape, node colors, labels, pie charts, or layouts may appear to be missing.

### Potential solution

Make sure the network was saved as a `.cys` session rather than as a network export or image export. Only the `.cys` format preserves the complete Cytoscape session, including layout, styles, labels, and chart overlays. Reopen the saved session immediately after saving to confirm that all settings persist.

## Resource availability

### Lead contact

Further information and requests for resources and reagents should be directed to and will be fulfilled by the lead contact, Alexander Aksenov.

### Technical contact

Technical questions on executing this protocol should be directed to and will be answered by the technical contact, Huang Lin.

### Materials availability

This study did not generate new materials.

### Data and code availability

- The HCC dataset was accessed through the Global Natural Products Social Molecular Networking platform (GNPS). Free registration with GNPS may be required to access these resources.
- MetVAE has been developed as a Python package and is available on PyPI. Code has been deposited to the GitHub repository and is publicly available as of the date of publication. Within the GitHub repository, the notebook `01_quickstart.ipynb` walks through and reproduces the example shown in the “Quickstart” section; `02_sim_study.ipynb` walks through and reproduces the simulation studies presented in the Supplementary Information and Figures S1 and S2; and `03_hcc.ipynb` walks through and reproduces the example shown in the “Step-by-step method details” section. Any additional information required to reanalyze the data reported in this paper is available from the lead contact upon request.

## Supplemental information index

### Supplementary Information PDF

Supplementary methods for the MetVAE framework, including model formulation, missing-data imputation, identifiability, sparsification, and simulation benchmarking results with Figures S1 and S2, related to Steps 8–10.

## Acknowledgments

This research of JTM was supported through funding by Department of Energy grant DE-SC002432 AA and AL were supported by the USDA NIFA GRANT 13665683. This research was supported in part by the Intramural Research Program of the NIH, National Institute of Environmental Health Sciences (ZIC ES103363). The contributions of the NIH author(s) are considered Works of the United States Government. The findings and conclusions presented in this paper are those of the author(s) and do not necessarily reflect the views of the NIH or the U.S. Department of Health and Human Services.

## Author contributions

JM and HL contributed equally to the theory and methodology described in this paper. AA and AL conducted metabolomics analyses. AK contributed to murine HCC study results interpretation. All numerical works and computations were conducted by HL, who developed MetVAE pipeline in Python that is freely and publicly available. AA and HL wrote the manuscript. All authors contributed to manuscript editing.

## Declaration of generative AI and AI-assisted technologies

During the preparation of this work, the author(s) used ChatGPT for grammatical review and language refinement. After using this tool or service, the author(s) reviewed and edited the content as needed and take(s) full responsibility for the content of the publication.

## References

1. Beger, R. D. (2013). A review of applications of metabolomics in cancer. Metabolites 3, 552–574.

2. Smolinska, A., Blanchet, L., Buydens, L. M., and Wijmenga, S. S. (2012). Nmr and pattern recognition methods in metabolomics: from data acquisition to biomarker discovery: a review. Analytica chimica acta 750, 82–97.

3. Koek, M. M., Jellema, R. H., van der Greef, J., Tas, A. C., and Hankemeier, T. (2011). Quantitative metabolomics based on gas chromatography mass spectrometry: status and perspectives. Metabolomics 7, 307–328.

4. Fenaille, F., Saint-Hilaire, P. B., Rousseau, K., and Junot, C. (2017). Data acquisition workflows in liquid chromatography coupled to high resolution mass spectrometry-based metabolomics: Where do we stand? Journal of Chromatography a 1526, 1–12.

5. Schrimpe-Rutledge, A. C., Codreanu, S. G., Sherrod, S. D., and McLean, J. A. (2016). Untargeted metabolomics strategies—challenges and emerging directions. Journal of the American Society for Mass Spectrometry 27, 1897–1905.

6. Aksenov, A. A., da Silva, R., Knight, R., Lopes, N. P., and Dorrestein, P. C. (2017). Global chemical analysis of biology by mass spectrometry. Nature Reviews Chemistry 1, 0054.

7. da Silva, R. R., Dorrestein, P. C., and Quinn, R. A. (2015). Illuminating the dark matter in metabolomics. Proceedings of the National Academy of Sciences 112, 12549–12550.

8. Wang, M., Carver, J. J., Phelan, V. V., Sanchez, L. M., Garg, N., Peng, Y., Nguyen, D. D., Watrous, J., Kapono, C. A., Luzzatto-Knaan, T. et al. (2016). Sharing and community curation of mass spectrometry data with global natural products social molecular networking. Nature biotechnology 34, 828–837.

9. Watrous, J., Roach, P., Alexandrov, T., Heath, B. S., Yang, J. Y., Kersten, R. D., Van Der Voort, M., Pogliano, K., Gross, H., Raaijmakers, J. M., et al. (2012). Mass spectral molecular networking of living microbial colonies. Proceedings of the National Academy of Sciences 109, E1743–E1752.

10. Nothias, L.-F., Petras, D., Schmid, R., Dü hrkop, K., Rainer, J., Sarvepalli, A., Protsyuk, I., Ernst, M., Tsugawa, H., Fleischauer, M., et al. (2020). Feature-based molecular networking in the gnps analysis environment. Nature methods 17, 905–908.

11. Coler, E. A., Melnik, A., Lotfi, A., Moradi, D., Ahiadu, B., Gomes, P. W. P., Patan, A., Dorrestein, P. C., Barnes, S., Boginski, V. et al. (2024). Ordering molecular diversity in untargeted metabolomics via molecular community networking. bioRxiv.

12. Aksenov, A. A., Laponogov, I., Zhang, Z., Doran, S. L., Belluomo, I., Veselkov, D., Bit-tremieux, W., Nothias, L. F., Nothias-Esposito, M., Maloney, K. N. et al. (2021). Auto-deconvolution and molecular networking of gas chromatography–mass spectrometry data. Nature biotechnology 39, 169–173.

13. Amara, A., Frainay, C., Jourdan, F., Naake, T., Neumann, S., Novoa-Del-Toro, E. M., Salek, R. M., Salzer, L., Scharfenberg, S., and Witting, M. (2022). Networks and graphs discovery in metabolomics data analysis and interpretation. Frontiers in Molecular Biosciences 9, 841373.

14. Liu, W., Guo, P., Dai, T., Shi, X., Shen, G., and Feng, J. (2021). Metabolic interactions and differences between coronary heart disease and diabetes mellitus: a pilot study on biomarker determination and pathogenesis. Journal of Proteome Research 20, 2364–2373.

15. Faubert, B., Solmonson, A., and DeBerardinis, R. J. (2020). Metabolic reprogramming and cancer progression. Science 368, eaaw5473.

16. Friedman, J., and Alm, E. J. (2012). Inferring correlation networks from genomic survey data. PLOS computational biology 8, e1002687.

17. Lin, H., Eggesbø, M., and Peddada, S. D. (2022). Linear and nonlinear correlation estimators unveil undescribed taxa interactions in microbiome data. Nature communications 13, 4946.

18. Kurtz, Z. D., Mü ller, C. L., Miraldi, E. R., Littman, D. R., Blaser, M. J., and Bonneau, R. A. (2015). Sparse and compositionally robust inference of microbial ecological networks. PLoS computational biology 11, e1004226.

19. Yoon, G., Gaynanova, I., and Mü ller, C. L. (2019). Microbial networks in spring-semi-parametric rank-based correlation and partial correlation estimation for quantitative microbiome data. Frontiers in genetics 10, 449195.

20. Bijlsma, S., Bobeldijk, I., Verheij, E. R., Ramaker, R., Kochhar, S., Macdonald, I. A., Van Ommen, B., and Smilde, A. K. (2006). Large-scale human metabolomics studies: a strategy for data (pre-) processing and validation. Analytical chemistry 78, 567–574.

21. Shalapour, S., Lin, X.-J., Bastian, I. N., Brain, J., Burt, A. D., Aksenov, A. A., Vrbanac, A. F., Li, W., Perkins, A., Matsutani, T. et al. (2017). Inflammation-induced iga+ cells dismantle anti-liver cancer immunity. Nature 551, 340–345.

22. Benjamini, Y., and Hochberg, Y. (1995). Controlling the false discovery rate: a practical and powerful approach to multiple testing. Journal of the Royal statistical society: series B (Methodological) 57, 289–300.

23. Cui, Y., Leng, C., and Sun, D. (2016). Sparse estimation of high-dimensional correlation matrices. Computational Statistics & Data Analysis 93, 390–403.

24. Best, C. A., and Laposata, M. (2003). Fatty acid ethyl esters: toxic non-oxidative metabolites of ethanol and markers of ethanol intake. Front Biosci 8, e202–17.

25. Singal, A. K. (2021). Similarities and differences between non-alcoholic fatty liver disease (nafld) & alcohol-associated liver disease (ald). Translational gastroenterology and hepatology 6, 1.

26. Díaz, L. A., Arab, J. P., Louvet, A., Bataller, R., and Arrese, M. (2023). The intersection between alcohol-related liver disease and nonalcoholic fatty liver disease. Nature Reviews Gastroenterology & Hepatology 20, 764–783.

27. Malnick, S. D., Alin, P., Somin, M., and Neuman, M. G. (2022). Fatty liver disease-alcoholic and non-alcoholic: similar but different. International Journal of Molecular Sciences 23, 16226.

