## Supplementary Information for "Protocol for constructing correlation-based molecular networks from large-scale untargeted metabolomics data"

#### Contents


1 MetVAE Methods 3

2 Simulation Studies 10

3 Supplementary Figures 13

### 1 MetVAE Methods

MetVAE introduces an integrated framework for correlation analysis in untargeted metabolomics: (1) Compositionality: it applies a centered log-ratio (CLR) transformation to account for the relative (compositional) nature of MS measurements; (2) Missing data: it uses VAE-based reconstructions to perform robust imputation while learning latent structure; (3) Covariate/confounder control: it incorporates covariates directly into the generative model to mitigate technical and biological confounding; and (4) High dimensionality: it employs Sparse Estimation of the Correlation (SEC) algorithm [1] to efficiently produce a sparse, positive-definite correlation matrix.

#### General workflow

Given a dataset of $N$ samples (indexed by $i$) and $D$ metabolites (indexed by $j$), we represent the log observed abundances of metabolites in sample $i$ as a vector $\mathbf{o}_i \in \mathbb{R}^D$, where $\mathbf{o}_i = [o_{ij}]_{j=1,\ldots,D}$. Some metabolites may be unobserved due to factors like random errors, stochastic fluctuations, data preprocessing limitations, or values below the limits of quantification (LOQ). This can result in zero entries. To denote the observation status, we employ the notation $(\cdot)^{\mathrm{o}}$ for the subvector of elements in positions where the abundances are observed (non-missing), and $(\cdot)^{\mathrm{m}}$ for the subvector of elements in positions where the abundances are missing. With these definitions in place, we outline the general workflow of the MetVAE model as follows:

| Missing Data Imputation |
|---|
| $\mathbf{y}_i^{\mathrm{o}} = \mathrm{clr}(\mathbf{o}_i^{\mathrm{o}})$<br>$\mathbf{y}_i = \iota\left(\mathbf{y}_i^{\mathrm{o}}\right)$ |
| Encoder |
| $p(\mathbf{z}_i) = \mathcal{N}(\mathbf{0}, \mathbf{I}_K)$<br>$q_{\boldsymbol{\phi}}(\mathbf{z}_i\|\mathbf{x}_i, \mathbf{y}_i) = \mathcal{N}\left(\mathbf{V}(\mathbf{y}_i - \mathbf{B}\mathbf{x}_i), \boldsymbol{\Omega}\right)$ |
| Decoder |
| $p_{\boldsymbol{\theta}}(\mathbf{y}_i\|\mathbf{z}_i, \mathbf{x}_i) = \mathcal{N}\left(\mathbf{W}\mathbf{z}_i + \mathbf{B}\mathbf{x}_i, \sigma^2\mathbf{I}_d\right)$ |
| Data Reconstruction |
| $\mathbf{y}_i^{\mathrm{m}} \sim p_{\boldsymbol{\theta}}(\mathbf{y}_i\|\mathbf{z}_i, \mathbf{x}_i)$<br>$\mathbf{y}_i = (\mathbf{y}_i^{\mathrm{o}}, \mathbf{y}_i^{\mathrm{m}})$ |

where

1. $\mathrm{clr}(\cdot)$: the CLR transformation. $\mathrm{clr}(\mathbf{x}) = \mathbf{x} - \bar{x}$ if $\mathbf{x}$ is in log scale,
2. $\mathbf{y} \in \mathbb{R}^D$: the CLR-transformed abundances,
3. $\iota\left(\mathbf{y}_i^{\mathrm{o}}\right)$ is an imputation function that transforms $\mathbf{y}_i^{\mathrm{o}}$ into a complete vector $\mathbf{y}_i$ such that $(\iota\left(\mathbf{y}_i^{\mathrm{o}}\right))^{\mathrm{o}} = \mathbf{y}_i^{\mathrm{o}}$,
4. $\mathbf{z}_i \in \mathbb{R}^K$: the latent variables, $K \leq D$,
5. $p(\mathbf{z}_i)$: the prior distribution of the latent variables,
6. $\mathbf{x}_i \in \mathbb{R}^P$: the covariates/confounders,

7. $\mathbf{B} \in \mathbb{R}^{D \times P}$: the effect sizes of covariates/confounders,

8. $q_{\boldsymbol{\phi}}(\mathbf{z}_i|\mathbf{x}_i, \mathbf{y}_i)$: the variational distribution represented by the encoder network, which is characterized by parameters $\boldsymbol{\phi} = (\mathbf{V}, \boldsymbol{\Omega})$,

9. $p_{\boldsymbol{\theta}}(\mathbf{y}_i|\mathbf{z}_i, \mathbf{x}_i)$: The posterior distribution represented by the decoder network, with parameters $\boldsymbol{\theta} = (\mathbf{W}, \sigma^2)$. Here, the weight matrix $\mathbf{W}$ is also known as the scaled principal components for the CLR-transformed data.

#### Objective function

In the standard VAE approach, we use the evidence lower bound (ELBO) as the objective function:

$$\mathcal{L}(\boldsymbol{\theta}, \boldsymbol{\phi}; y, x) = \frac{1}{N} \sum_{i=1}^{N} \mathcal{L}(\boldsymbol{\theta}, \boldsymbol{\phi}; \mathbf{y}_i, \mathbf{x}_i),$$

where

$$\begin{aligned}
\mathcal{L}(\boldsymbol{\theta}, \boldsymbol{\phi}; \mathbf{y}_i, \mathbf{x}_i) &= \mathbb{E}_{\mathbf{z}_i \sim q_{\boldsymbol{\phi}}(\mathbf{z}_i|\mathbf{x}_i, \mathbf{y}_i)} \left[ \ln \frac{p_{\boldsymbol{\theta}}(\mathbf{y}_i|\mathbf{z}_i, \mathbf{x}_i) p(\mathbf{z}_i)}{q_{\boldsymbol{\phi}}(\mathbf{z}_i|\mathbf{x}_i, \mathbf{y}_i)} \right] \\
&= \mathbb{E}_{\mathbf{z}_i \sim q_{\boldsymbol{\phi}}(\mathbf{z}_i|\mathbf{x}_i, \mathbf{y}_i)} \left[ \ln \frac{p_{\boldsymbol{\theta}}(\mathbf{y}_i, \mathbf{z}_i|\mathbf{x}_i)}{q_{\boldsymbol{\phi}}(\mathbf{z}_i|\mathbf{x}_i, \mathbf{y}_i)} \right] \\
&\leq \ln \mathbb{E}_{\mathbf{z}_i \sim q_{\boldsymbol{\phi}}(\mathbf{z}_i|\mathbf{x}_i, \mathbf{y}_i)} \left[ \frac{p_{\boldsymbol{\theta}}(\mathbf{y}_i, \mathbf{z}_i|\mathbf{x}_i)}{q_{\boldsymbol{\phi}}(\mathbf{z}_i|\mathbf{x}_i, \mathbf{y}_i)} \right] \quad \text{by Jensen's Inequality} \\
&= \ln p_\theta \left( \mathbf{y}_i \mid \mathbf{x}_i \right).
\end{aligned}$$

Alternatively, for a tighter bound on the log-likelihood, one can also use the Importance Weighted Autoencoder (IWAE) [2]. While sharing the same architecture as a VAE, the IWAE introduces multiple samples during the optimization process, leading to an improved tightness of the bound. The objective function is:

$$\mathcal{L}_M(\boldsymbol{\theta}, \boldsymbol{\phi}; y, x) = \sum_{i=1}^{N} \mathbb{E} \left[ \log \frac{1}{M} \sum_{m=1}^{M} w_{mi} \right],$$

where $M$ refers to the number of importance samples used to estimate the variational lower bound, and for $m \leq M$ and $i \leq N$,

$$w_{mi} = \frac{p_{\boldsymbol{\theta}} \left( \mathbf{y}_{mi} \mid \mathbf{z}_{mi}, \mathbf{x}_i \right) p \left( \mathbf{z}_{mi} \right)}{q_{\boldsymbol{\phi}} \left( \mathbf{z}_{mi} \mid \mathbf{x}_i, \mathbf{y}_{mi} \right)},$$

and $\left( \mathbf{z}_{1i}, \mathbf{y}_{1i} \right), \ldots, \left( \mathbf{z}_{Mi}, \mathbf{y}_{Mi} \right)$ are $M$ i.i.d. samples from $q_{\boldsymbol{\phi}} \left( \mathbf{z}_i \mid \mathbf{y}_i, \mathbf{x}_i \right) p_{\boldsymbol{\theta}} \left( \mathbf{y}_i \mid \mathbf{z}_i, \mathbf{x}_i \right)$.

#### Missing data imputation

During the training phase, at each epoch, missing values for each metabolite are imputed using randomly generated values from a censored Normal distribution. This approach diverges significantly from the method of adding pseudo-counts to the original data, which can introduce artificial correlations. A key advantage of our imputation strategy is its capacity to mimic the distribution of the complete underlying data, effectively reducing the occurrence of false positives. Moreover, by imputing missing values with different values in each epoch, the strategy encourages the VAE model to de-emphasize missing data, instead prioritizing the analysis of non-missing data. This approach aids the model in accurately learning the correct low-dimensional latent representations, culminating in a more accurate reconstruction of the data.

For simplicity, we will omit the metabolite index $j$ in further discussions, as the imputation method is applied individually to each metabolite. Assuming the complete CLR-transformed abundance $y_i$ follows a Normal distribution $\mathcal{N}(\mu, \sigma^2)$, where $\mu = \boldsymbol{\beta}^T \mathbf{x}_i$ and $\mathbf{x}_i$ encompasses the intercept term along with any covariates or confounders. Setting a limit of quantification (LOQ) represented by $\tau$, the observed values are then assumed to follow a censored Normal distribution, expressed as:

$$y_i = \begin{cases} y_i & \text{if } y_i \geq \tau, \\ 0 & \text{otherwise.} \end{cases}$$

The log-likelihood function is:

$$\begin{aligned} \ell(\boldsymbol{\beta}, \sigma^2) &= \sum_{i:y_i \geq \tau} \left\{ -\frac{1}{2} \ln 2\pi - \ln \sigma - \frac{(y_i - \mu_i)^2}{2\sigma^2} \right\} + \sum_{i:y_i < \tau} \ln \Phi \left( \frac{\tau - \mu_i}{\sigma} \right) \\ &\propto - \ln \sigma \cdot \left( (\mathbf{1}^{\mathrm{o}})^T \mathbf{1}^{\mathrm{o}} \right) - \frac{1}{2\sigma^2} \left( (\mathbf{y} - \boldsymbol{\mu})^{\mathrm{o}} \right)^T (\mathbf{y} - \boldsymbol{\mu})^{\mathrm{o}} + \left( \left( \ln \Phi \left( \frac{\tau - \mu_i}{\sigma} \right) \right)^{\mathrm{m}} \right)^T \mathbf{1}^{\mathrm{m}}. \end{aligned}$$

And the corresponding gradients for $\boldsymbol{\beta}$ and $\sigma$ are:

$$\begin{aligned} \frac{\partial \ell}{\partial \boldsymbol{\beta}} &= \sum_{i:y_i \geq \tau} \frac{(y_i - \mu_i)^2}{\sigma^2} \mathbf{x}_i - \sum_{i:y_i < \tau} \frac{\phi\left(\frac{\tau - \mu_i}{\sigma}\right)}{\sigma \Phi\left(\frac{\tau - \mu_i}{\sigma}\right)} \mathbf{x}_i \\ &= \frac{1}{\sigma^2} (\mathrm{x}^{\mathrm{o}})^T (\mathbf{y} - \mathrm{x}\boldsymbol{\beta})^{\mathrm{o}} - \frac{1}{\sigma} (\mathrm{x}^{\mathrm{m}})^T \left( \frac{\phi\left(\frac{\tau - \boldsymbol{\mu}}{\sigma}\right)}{\sigma \Phi\left(\frac{\tau - \boldsymbol{\mu}}{\sigma}\right)} \right)^{\mathrm{m}}, \end{aligned}$$

$$\begin{aligned} \frac{\partial \ell}{\partial \sigma} &= \sum_{i:y_i \geq \tau} \left\{ -1 + \frac{(y_i - \mu_i)^2}{\sigma^2} \right\} - \sum_{i:y_i < \tau} \frac{\phi\left(\frac{\tau - \mu_i}{\sigma}\right)}{\Phi\left(\frac{\tau - \mu_i}{\sigma}\right)} \frac{\tau - \mu_i}{\sigma} \\ &= -(\mathbf{1}^{\mathrm{o}})^T \mathbf{1}^{\mathrm{o}} + \frac{1}{\sigma^2} \left( (\mathbf{y} - \boldsymbol{\mu})^{\mathrm{o}} \right)^T (\mathbf{y} - \boldsymbol{\mu})^{\mathrm{o}} - \frac{1}{\sigma} \left( \left( \frac{\phi\left(\frac{\tau - \boldsymbol{\mu}}{\sigma}\right)}{\Phi\left(\frac{\tau - \boldsymbol{\mu}}{\sigma}\right)} \right)^{\mathrm{m}} \right)^T (\tau - \boldsymbol{\mu})^{\mathrm{m}}, \end{aligned}$$

where

(1) $\phi$ and $\Phi$ represent the probability density function (pdf) and cumulative distribution function (cdf) for the standard Normal distribution, respectively,

(2) $\mathbf{x}$, an $N \times P$ matrix, denotes the covariates (including the intercept term) for all samples.

The estimates of $\boldsymbol{\beta}$ and $\sigma$ can be obtained by maximizing the log-likelihood function. This optimization can be efficiently conducted using algorithms such as the Broyden–Fletcher–Goldfarb–Shanno (BFGS) algorithm [3]. For enhanced numerical stability, it is advisable to compute gradients with respect to $\ln \sigma$ rather than $\sigma$ directly. The gradient for $\sigma$ is straightforward to calculate once it is recognized that $\frac{\partial \sigma}{\partial \ln \sigma} = \sigma$.

During the inference phase, we implement multiple imputation by utilizing the reconstructed values from the VAE decoder. Specifically, observed (non-missing) values are retained, while missing entries are imputed with their VAE-reconstructed counterparts in each iteration. Final pairwise correlation estimates are computed by averaging correlation matrices across all imputed datasets, thereby accounting for imputation uncertainty.

#### Identifiability of the correlation

We firstly assume the following data-generating process for the log observed metabolic abundances, which is similar to the model proposed in the Analysis of Compositions of Microbiomes with Bias Correction 2 (ANCOM-BC2) [4].

**Assumption 1** (Data-generating process). *The log-transformed observed abundance, $o_{ij}$, can be decomposed additively into the following components::*

$$o_{ij} = s_i + c_j + a_{ij} + e_{ij},$$

*where $s_i$ represents the sample-specific bias, $c_j$ represents the feature-specific bias, $a_{ij}$ is the log absolute abundance, and $e_{ij}$ is the random error, with $e_{ij} \sim_{i.i.d.} F_e$ such that $\mathrm{E}(e_{ij}) = 0, \mathrm{Var}(e_{ij}) = \sigma_e^2$.*

*Furthermore, it is assumed that the log absolute abundance is independent of the random error:*

$$a_{ij} \perp e_{ij}.$$

Note that in the presence of covariates or confounders, the log absolute abundance, $a_{ij}$ can be further decomposed into $a_{ij} = \mathbf{b_j}^T \mathbf{x_i} + \epsilon_{ij}$, where $\mathbf{b_j}$ represents the effect of the covariates $\mathbf{x_i}$ on the log absolute abundance, and $\epsilon_{ij}$ is the residual. Consequently, the data-generating model can be readily extended as follows:

$$\begin{aligned} &o_{ij} = s_i + c_j + \mathbf{b_j}^T \mathbf{x_i} + \epsilon_{ij} + e_{ij}, \\ &\epsilon_{ij} \perp e_{ij}. \end{aligned}$$

To avoid redundancy, we will focus solely on the case where covariates or confounders are absent in the following discussion.

Next, we define the true covariance matrix, $\Sigma_0 = \left(\sigma_{lm}^0\right)_{D \times D}$ as follows:

$$\sigma_{lm}^0 = \mathrm{Cov}(a_l, a_m),$$

where $a_l, a_m$ are log absolute abundances for feature $l$ and $m$ (with $l, m \in \{1, \ldots, D\}$), respectively. We assume that $\Sigma_0$ belongs to a class of sparse covariance matrices [5].

**Assumption 2** (Sparse covariance matrices).

$$\mathcal{U}(q, K) = \left\{ \Sigma : \Sigma \succ 0, \max_j \sigma_{jj} \leq K, \max_l \sum_{m=1}^{D} |\sigma_{lm}|^q \leq o(D) \right\},$$

*where $0 \leq q < 1$, $\Sigma \succ 0$ denotes that $\Sigma$ is positive definite, and $o(\cdot)$ is the little-o notation which means "is ultimately smaller than".*

This assumption delineates the covariance matrix for log absolute abundances as sparse, which is a common assumption in the estimation of high-dimensional covariance matrices [6, 7, 8, 9].

Lastly, we make the following assumption regarding the noise-to-signal ratio.

**Assumption 3** (Small noise-to-signal ratio).

$$\sigma_e^2 \ll \sigma_{jj}^0.$$

With all the above assumptions in place, we present the following theorem regarding the identifiability of the true correlations between the log absolute abundances.

**Theorem 1.1.** *Let the true Pearson correlation coefficient matrix, calculated based on the log absolute abundances,* $a_j$*, be denoted as* $R_0 = \left(\rho^0_{lm}\right)_{D\times D}$*, where:*

$$\rho^0_{lm} = \frac{\sigma^0_{lm}}{\sqrt{\sigma^0_{ll}\sigma^0_{mm}}}.$$

*Additionally, define covariance matrix for the CLR-transformed abundances,* $y_j$*, and the matrix of Variance of Log-Ratios (VLRs) as* $\Sigma = (\sigma_{lm})_{D\times D}$ *and* $T = (t_{lm})_{D\times D}$*, respectively, where:*

$$\sigma_{lm} = \mathrm{Cov}(y_l - y_m),$$
$$t_{lm} = \mathrm{Var}(o_l - o_m).$$

*Given Assumptions 1, 2, and 3, the following estimator:*

$$\rho_{lm} = \frac{\sigma_{ll} + \sigma_{mm} - t_{lm}}{2\sqrt{\sigma_{ll}\sigma_{mm}}}, \tag{1}$$

*asymptotically approximates* $\rho^0_{lm}$*, i.e.,*

$$\rho_{lm} \to \rho^0_{lm} \quad \text{as } D \to \infty. \tag{2}$$

*Proof:* Starting with the definition of VLR, we express $t_{lm}$ as follows:

$$\begin{aligned} t_{lm} &= \mathrm{Var}(o_l - o_m) = \mathrm{Var}(c_l + a_l + e_l - c_m - a_m - e_m) = \mathrm{Var}(a_l - a_m) + \mathrm{Var}(e_l - e_m), \\ &= \sigma^0_{ll} + \sigma^0_{mm} - 2\sigma^0_{lm} + 2\sigma^2_e = \sigma^0_{ll} + \sigma^0_{mm} - 2\rho^0_{lm}\sqrt{\sigma^0_{ll}\sigma^0_{mm}} + 2\sigma^2_e. \end{aligned} \tag{3}$$

Rearranging equation (3) yields that:

$$\rho^0_{lm} = \frac{\sigma^0_{ll} + \sigma^0_{mm} + 2\sigma^2_e - t_{lm}}{2\sqrt{\sigma^0_{ll}\sigma^0_{mm}}}. \tag{4}$$

To demonstrate that $\rho_{lm}$ asymptotically approximates $\rho^0_{lm}$, we begin by showing that numerator of $\rho_{lm}$ is asymptotically indistinguishable from the numerator of $\rho^0_{lm}$. Based on Assumption 1 and the definition of the CLR transformation, we have:

$$\begin{aligned} o_j &= s + c_j + a_j + e_j, \\ y_j &= o_j - \frac{1}{D}\sum_{j=1}^{D} o_j = c_j - \bar{c} + a_j - \bar{a} + e_j - \bar{e}. \end{aligned}$$

Thus,

$$\begin{aligned} \mathrm{Var}(y_j) &= \mathrm{Var}(c_j - \bar{c} + a_j - \bar{a} + e_j - \bar{e}) \\ &= \underbrace{\mathrm{Var}(a_j - \bar{a})}_{①} + \underbrace{\mathrm{Var}(e_j - \bar{e})}_{②}. \end{aligned}$$

It is straightforward to verify that, for ②:

$$② = \sigma^2_e + \frac{1}{D^2}D\sigma^2_e - \frac{2}{D}\sigma^2_e = \frac{D-1}{D}\sigma^2_e \to \sigma^2_e \quad \text{as } D \to \infty. \tag{5}$$

For ①, we have:

$$① = \sigma^0_{jj} - \frac{2}{D}\sum_{j'}\sigma^0_{jj'} + \frac{1}{D^2}\sum_{j}\sigma^0_{jj} + \frac{1}{D^2}\sum_{j}\sum_{j'\neq j}\sigma^0_{jj'} = \frac{D-2}{D}\sigma^0_{jj} + \frac{1}{D^2}\sum_{j}\sigma^0_{jj} - \frac{2}{D}\sum_{j'\neq j}\sigma^0_{jj'} + \frac{1}{D^2}\sum_{j}\sum_{j'\neq j}\sigma^0_{jj'}. \tag{6}$$

Based on Assumption 2, one can show that:

$$\sum_{j'\neq j}\sigma^0_{jj'} \leq \sum_{j'}\sigma^0_{jj'} \leq \sum_{j'}|\sigma^0_{jj'}| \leq \max_j \sum_{j'}|\sigma^0_{jj'}|^{1-q}|\sigma^0_{jj'}|^q \leq \max_j \sum_{j'}(\sigma^0_{jj}\sigma^0_{j'j'})^{(1-q)/2}|\sigma^0_{jj'}|^q \leq K^{1-q}o(D) = o(D).$$

Similarly, we obtain:

$$\sum_{j}\sum_{j'\neq j}\sigma^0_{jj'} \leq \sum_{j}\sum_{j'}\sigma^0_{jj'} \leq D\max_{j'}\sum_{j'}\sigma^0_{jj'} = o(D^2).$$

Therefore, we have:

$$① \to \sigma^0_{jj}, \quad \text{as } D \to \infty.$$

Combining ① and ②, we demonstrate that:

$$\sigma_{jj} := \mathrm{Var}(y_j) \to \sigma^0_{jj} + \sigma^2_e \quad \text{as } D \to \infty.$$

Thus, the numerator of $\rho_{lm}$,

$$\sigma_{ll} + \sigma_{mm} - t_{lm} \to \sigma^0_{ll} + \sigma^0_{mm} + 2\sigma^2_e - t_{lm}, \quad \text{as } D \to \infty,$$

converges to the numerator of $\rho^0_{lm}$.

Next, we demonstration the approximation of the denominators between $\rho_{lm}$ and $\rho^0_{lm}$. Based on Assumption 3, we have:

$$\sigma_{jj} \to \sigma^0_{jj} + \sigma^2_e \approx \sigma^0_{jj} \quad \text{as } D \to \infty.$$

Therefore, the denominator of $\rho_{lm}$:

$$2\sqrt{\sigma_{ll}\sigma_{mm}} \to 2\sqrt{(\sigma^0_{ll} + \sigma^2_e)(\sigma^0_{mm} + \sigma^2_e)} \approx 2\sqrt{\sigma^0_{ll}\sigma^0_{mm}} \quad \text{as } D \to \infty,$$

approximates the denominator of $\rho^0_{lm}$.

Combining all the above demonstrations, we have:

$$\rho_{lm} \approx \rho^0_{lm}, \quad \text{as } D \to \infty.$$

VLR can be effectively calculated utilizing the CLR-transformed data. Consequently, Theorem 3 establishes a connection between the correlations among log absolute abundances and their CLR-transformed counterparts.

*Remark* 1.2. With a limited number of features, i.e., when $D$ is not large, we suggest the following modifications. Based on equations (5) and (6), and under Assumption 2, which allows us to "ignore" all covariances, we have:

$$\sigma_{jj} = \frac{D-2}{D}\sigma^0_{jj} + \frac{1}{D^2}\sum_{j}\sigma^0_{jj} + \frac{D-1}{D}\sigma^2_e. \tag{7}$$

With a simple rearrangement of equation (7), we obtain:

$$\sigma_{jj}^0 + \sigma_e^2 = \frac{D}{D-2}\sigma_{jj} - \frac{1}{D(D-2)}\sum_j \sigma_{jj}^0 - \frac{1}{D-2}\sigma_e^2. \tag{8}$$

On the other hand, summing equation (7) across features, $j$, leads to:

$$\sum_j \sigma_{jj} = \frac{D-1}{D}\sum_j \sigma_{jj}^0 + (D-1)\sigma_e^2. \tag{9}$$

Substituting equation (9) into equation (8) leads to the modified estimator:

$$\sigma_{jj}^* = \frac{D}{D-2}\sigma_{jj} - \frac{1}{(D-1)(D-2)}\sum_j \sigma_{jj}, \tag{10}$$

and

$$\rho_{lm}^* = \frac{\sigma_{ll}^* + \sigma_{mm}^* - t_{lm}}{2\sqrt{\sigma_{ll}^*\sigma_{mm}^*}}. \tag{11}$$

#### Sparse correlation estimation

Let $R_n$ denote the empirical correlation matrix estimated from the CLR-transformed abundances, $y_j$. We estimate the sparse correlation matrix using the Sparse Estimation of the Correlation matrix (SEC) algorithm [1]:

$$\hat{R} = \arg\min_R \frac{1}{2}\|R - R_n\|_F^2 + \lambda|R|_1, \text{ s.t. } R \succeq \epsilon I, \sigma_{jj} = 1, j = 1, \dots, D. \tag{12}$$

where $\|\cdot\|$ is the Frobenius norm, $R \succeq \epsilon I$ indicates that $R - \epsilon I$ is positive semidefinite, $\epsilon$ is set to $10^{-5}$ by default, and $\lambda$ is the regularization parameter for the $l_1$ norm.

When the distributions of CLR-transformed abundances, $y_j$, exhibit either exponential-type or polynomial-type tail probabilities [1], the convergence of $\hat{R}$ to $R$ in Frobenius norm is established in the following theorem.

**Theorem 1.3.** *Denote the non-diagonal support of $R$ as $A_0 = \{l, m : l \neq m, \rho_{lm} \neq 0\}$ and its cardinality as $q$. If we set $\lambda = K\sqrt{\frac{\log D}{N}}$ for some constant $K$, then:*

$$\|\hat{R} - R\|_F^2 = O_p(q\frac{\log D}{N}).$$

The proof of this theorem largely follows the proof of Theorem 2 provided in the original SEC paper. Interested readers are encouraged to refer to that paper for detailed explanations.

In practice, the tuning parameter $\lambda$ is selected in a data-driven way by $K$-fold cross-validation. Concretely, for each candidate value of $\lambda$ in a grid, we repeatedly: (i) fit SEC on the training folds to obtain $\hat{R}^{(-k)}(\lambda)$, and (ii) evaluate its discrepancy to the empirical correlation matrix on the held-out fold, $R_n^{(k)}$, using the Frobenius loss $\|\hat{R}^{(-k)}(\lambda) - R_n^{(k)}\|_F^2$. The final choice $\hat{\lambda}$ minimizes the average validation loss across folds (ties favoring smaller $\lambda$). This cross-validated $\hat{\lambda}$ is then used to refit SEC on the full data.

As a simpler, testing-based alternative to SEC, we also consider a sparsification procedure based on pairwise $p$-values. For each pair $(l, m)$ with $l < m$, we test

$$H_{0,lm} : \rho_{lm} = 0 \quad \text{vs.} \quad H_{A,lm} : \rho_{lm} \neq 0$$

using a Fisher $z$-transformation of the empirical correlation $\rho_{lm}$ and obtain a $p$-value $p_{lm}$. We then apply a multiple-testing correction, such as the Benjamini–Hochberg (BH) procedure [10], to the collection $\{p_{lm} : 1 \leq l < m \leq D\}$ at a target FDR level $\alpha$, and construct a sparse estimator by setting

$$\hat{\rho}_{lm} = \begin{cases} \rho_{lm}, & \text{if } \tilde{p}_{lm} \leq \alpha, \\ 0, & \text{otherwise,} \end{cases} \tag{13}$$

where $\tilde{p}_{lm}$ denotes the adjusted $p$-value for edge $(l, m)$, and symmetry is enforced by defining $\hat{\rho}_{ml} = \hat{\rho}_{lm}$.

Theoretical FDR guarantees for BH rely on assumptions such as independence or certain forms of positive regression dependence among the $p$-values. These assumptions are not strictly satisfied in our setting, where the CLR-transformed abundances induce strong and complex dependencies among the pairwise test statistics, and the pairwise testing framework itself creates a substantial amount of dependence among the test statistics as many tests share common variables and are computed from the same finite sample. Consequently, the BH-adjusted $p$-values should be viewed as an exploratory device rather than a procedure with formal FDR control. In simulations (data not shown), the SEC-based estimator tends to exhibit better empirical FDR control, whereas the p-value filtering approach generally achieves higher power (detects more true edges) at the cost of some inflation in FDR.

From a practical perspective, the two sparsification strategies are complementary. The SEC estimator produces a positive semidefinite correlation matrix with unit diagonal by construction, and it borrows strength across all entries of $R$ through the global $\ell_1$ penalty; this yields a coherent, regularized network estimate that is well suited for downstream tasks that require a valid covariance or correlation matrix. In contrast, the $p$-value filtering procedure is conceptually simple, computationally cheap, and easily interpretable on an edge-by-edge basis, making it attractive as an exploratory tool for screening potential associations. In applications where strict control of false discoveries and a well-posed correlation matrix are critical, we recommend using SEC as the primary estimator, with the $p$-value filtered network used as a complementary, higher-power but less rigorously calibrated exploratory summary.

### 2 Simulation Studies

#### Simulation details

**Simulation design and evaluation metrics.** We evaluated MetVAE's performance on simulated data with known "ground truth", enabling rigorous assessment of true positive rate (TPR) and false discovery rate (FDR) under controlled conditions that recapitulate key challenges of untargeted metabolomics. We compared MetVAE to widely used alternatives, including Pearson correlation coefficient, SECOM [9], SparCC [7], SPIEC-EASI [11], and SPRING [12]. MetVAE produced a sparse correlation matrix using the SEC algorithm [1]. For competing methods with built-in sparsity mechanisms, we used their recommended/default procedures. For SPIEC-EASI and SPRING, we used neighborhood selection (MB) [13], which has been reported to outperform graphical lasso in numerical experiments [12, 14]. For SECOM (Thresholding), we used the default hard-thresholding implementation. For methods that output $P$-values but do not directly enforce sparsity (Pearson, SparCC, and SECOM (Filtering)), we constructed sparse matrices by retaining correlations whose Benjamini–Hochberg (BH)-adjusted $P$-values were below 0.05 [10]. Because pairwise correlation tests are not independent and may violate assumptions underlying BH, we treat this filtering primarily as a pragmatic, comparable baseline rather than a guarantee of strict FDR control.

**Data generation** Our simulations began by defining the number of samples ($N$) and features ($D$), along with the number of correlated feature pairs ($cor\_pairs$). We explored three sample-to-feature ($N/D$) ratios to reflect different dimensional settings: a low-dimensional case with $N = 100, D = 50$, a high-dimensional scenario with $N = 100, D = 200$, and an extreme high-dimensional case with $N = 100, D = 500$. For each scenario, we predetermined that $0.2D$ pairs of features (or $0.4D$ individual features) should exhibit true correlations. Each feature, representing a metabolite, was modeled with a true absolute abundance profile following a log-normal distribution (Lognormal($\mu, \sigma^2$)), where $\mu \in [10, 14]$ and $\sigma = 1$. We created a correlation matrix starting as an identity matrix and then modified non-diagonal elements to establish predefined correlations ranging from $-0.7$ to $0.7$, ensuring a balance of both strong negative and positive correlations. To generate observed abundance, biases were incorporated into the absolute abundances as follows: Each sample was assigned a unique bias, randomly drawn from a uniform distribution ranging between $1 \times 10^{-3}$ and $1 \times 10^{-1}$, and each feature received a distinct bias from a separate uniform distribution ranging between $1 \times 10^{-1}$ and 1. These biases were logarithmically transformed and then added respectively to the log-transformed absolute abundance data.

**Addition of covariates** To mirror real-world complexities, both continuous and categorical covariates were integrated into our simulations. These covariates, extracted from provided metadata ($x$), were incorporated using lists of continuous ($cont\_list$) and categorical ($cat\_list$) variables, with the latter encoded via one-hot encoding for model inclusion. The influence of these covariates on metabolite abundances was quantified by modeling log fold-change ($es$), where $es$ varied between $-2$ and 2. 10% ($da\_prop = 0.1$) of the features were randomly affected.

**Inclusion of missing data** We varied the proportion of missing values ($zero\_prop$) from 0% (representing complete data) to 30%, simulating missing data primarily due to values falling below the limits of quantification (LOQ). For each feature, abundances that were less than the ($zero\_prop$ quantile of all generated abundances were set to zero, reflecting a missing not at random (MNAR) mechanism.

#### Simulation results

**Benchmark under compositionality and missingness.** We first evaluated performance under compositionality with increasing levels of missingness in the absence of confounders (Figure S1). Pearson correlations performed poorly on compositional data, yielding severely inflated FDR when missingness was low. For example, in a low-dimensional setting ($N = 100$, $D = 50$) with no missing values, Pearson achieved an FDR close to 1, indicating that most detected edges were spurious. As missingness increased, Pearson (computed from complete observations) showed marked loss of TPR, reflecting reduced effective sample size. Among the conditional-dependence approaches, SPIEC-EASI and SPRING exhibited weak control of false positives, with high FDR across scenarios. In the low-dimensional setting ($N = 100$, $D = 50$), SPRING reached an average FDR of ∼85%, approaching 100% in high-dimensional settings; its TPR also dropped substantially, including to ∼50% for $N = 100$, $D = 200$. SPIEC-EASI achieved higher TPR than SPRING in most settings but still showed elevated FDR, increasing from ∼40% in the low-dimensional setting to >80% in high-dimensional settings. Among correlation-based methods, SparCC showed the highest FDR (averaging $\sim$50% in the low-dimensional setting) with TPR comparable to SECOM (Thresholding) and SECOM (Filtering). However, SparCC's permutation-based $P$-value estimation was computationally prohibitive in high dimensions (e.g., $\sim$6 hours per dataset for $N = 100$, $D = 200$ on a single CPU), and we therefore did not include SparCC in subsequent high-dimensional benchmarks. SECOM (Thresholding) achieved the lowest FDR overall, at the cost of a modest reduction in TPR relative to SECOM (Filtering), consistent with prior reports [9]. MetVAE matched the low FDR of SECOM (Thresholding) while maintaining consistently high TPR across all conditions. Notably, MetVAE's TPR remained near or above 90% and exceeded both SECOM variants when missingness increased (e.g., at 30% missingness), highlighting improved robustness when complete-case approaches become underpowered.

**Benchmark under compositionality, missingness, and confounding.** Real metabolomics studies often include technical and biological confounders (e.g., batch effects, diet, age). We therefore extended simulations by introducing one continuous and one binary confounder. Results are summarized in Figure S2. As expected, methods that do not model confounding exhibited inflated FDR and reduced TPR across scenarios. In contrast, because MetVAE explicitly incorporates confounders in its generative model, it maintained FDR at levels comparable to the no-confounder setting and substantially below all competing methods, while preserving the highest TPR across dimensionality and missingness regimes. These results indicate that MetVAE can recover correlation structure more faithfully in settings that more closely resemble large-scale metabolomics studies.

### 3 Supplementary Figures

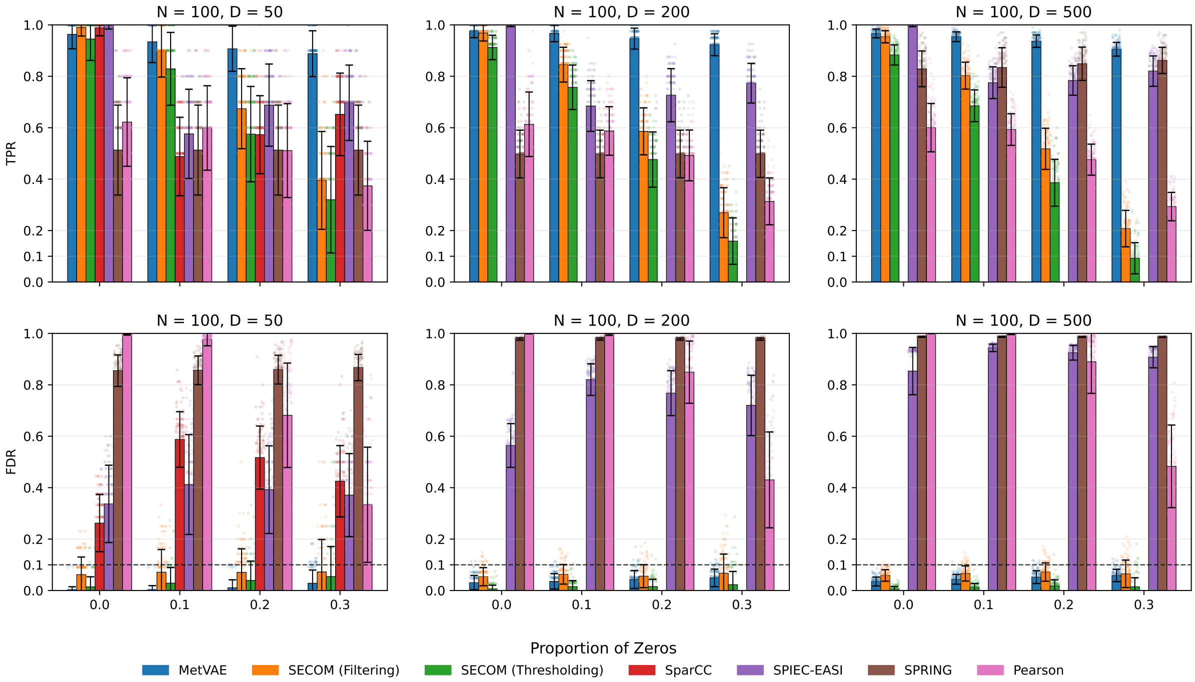


**Figure S1. Comparison of methods for detecting correlations under compositionality and missingness in the absence of confounders**. True absolute abundance data were generated from log-normal distributions, with observed abundances derived by adding sample-specific sampling fractions and taxon-specific sequencing efficiencies to the absolute abundances. The columns across the figure represent different simulation settings defined by combinations of sample size ($N$) and number of features ($D$). The x-axis indicates the proportion of missing data, ranging from 0 to 0.3, while the y-axis depicts both the true positive rate (TPR) and the false discovery rate (FDR), which are illustrated in the upper and lower panels of the figure, respectively. The results are summarized as the mean values of FDR and TPR, accompanied by standard deviations represented as error bars, derived from 100 simulation iterations for each specific $N/D$ configuration. Data points are overlaid on the bar charts as dots with jittering effects. The color scheme and labels identifying each method are displayed at the top of the graph.

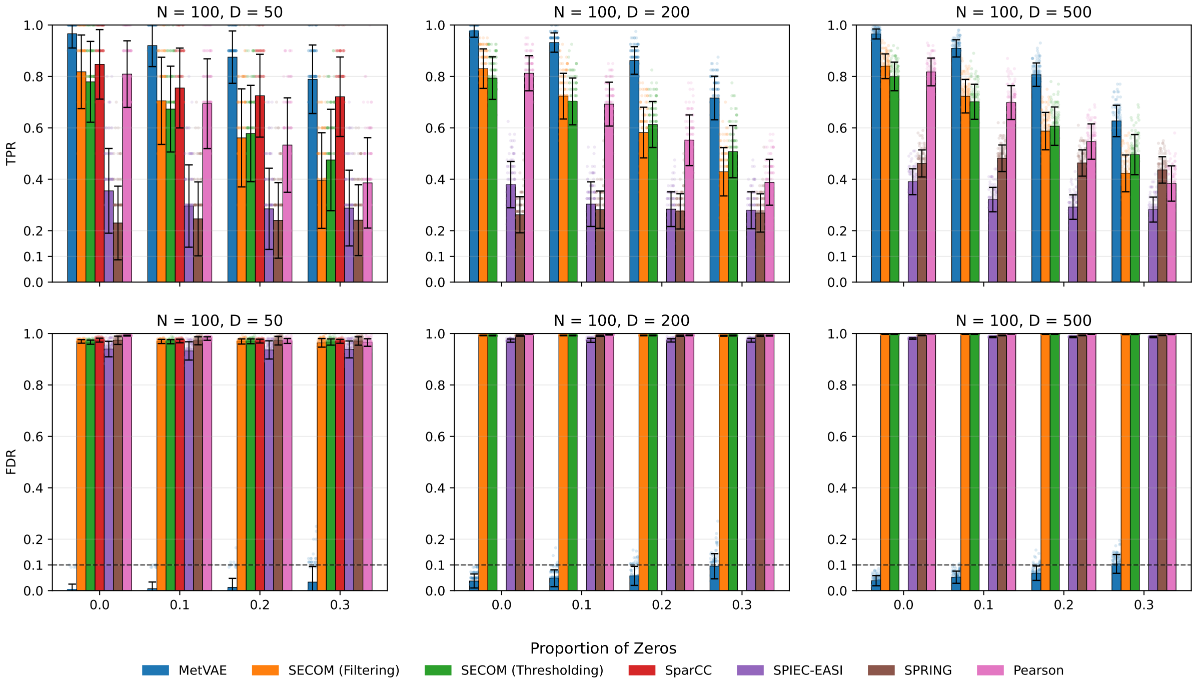


**Figure S2. Comparison of methods for detecting correlations under compositionality, missingness, and confounding**. True absolute abundance data were generated from log-normal distributions, incorporating effect sizes of one continuous and one binary confounder into the abundance profiles of differentially abundant metabolites. The observed abundances were derived by adding sample-specific sampling fractions and taxon-specific sequencing efficiencies to the absolute abundances. The columns across the figure represent different simulation settings defined by combinations of sample size ($N$) and number of features ($D$). The x-axis indicates the proportion of missing data, ranging from 0 to 0.3, while the y-axis depicts both the true positive rate (TPR) and the false discovery rate (FDR), which are illustrated in the upper and lower panels of the figure, respectively. The results are summarized as the mean values of FDR and TPR, accompanied by standard deviations represented as error bars, derived from 100 simulation iterations for each specific $N/D$ configuration. Data points are overlaid on the bar charts as dots with jittering effects. The color scheme and labels identifying each method are displayed at the top of the graph.

### References


[1] Ying Cui, Chenlei Leng, and Defeng Sun. Sparse estimation of high-dimensional correlation matrices. *Computational Statistics & Data Analysis*, 93:390–403, 2016.

[2] Yuri Burda, Roger Grosse, and Ruslan Salakhutdinov. Importance weighted autoencoders. *arXiv preprint arXiv:1509.00519*, 2015.

[3] Roger Fletcher. *Practical methods of optimization*. John Wiley & Sons, 2000.

[4] Huang Lin and Shyamal Das Peddada. Multigroup analysis of compositions of microbiomes with covariate adjustments and repeated measures. *Nature Methods*, 21(1):83–91, 2024.

[5] Peter J Bickel and Elizaveta Levina. Covariance regularization by thresholding. *The Annals of Statistics*, 36(6):2577–2604, 2008.

[6] Adam J Rothman, Elizaveta Levina, and Ji Zhu. Generalized thresholding of large covariance matrices. *Journal of the American Statistical Association*, 104(485):177–186, 2009.

[7] Jonathan Friedman and Eric J Alm. Inferring correlation networks from genomic survey data. *PLOS computational biology*, 8:e1002687, 2012.

[8] Yuanpei Cao, Wei Lin, and Hongzhe Li. Large covariance estimation for compositional data via composition-adjusted thresholding. *Journal of the American Statistical Association*, 114(526):759–772, 2019.

[9] Huang Lin, Merete Eggesbø, and Shyamal Das Peddada. Linear and nonlinear correlation estimators unveil undescribed taxa interactions in microbiome data. *Nature communications*, 13(1):4946, 2022.

[10] Yoav Benjamini and Yosef Hochberg. Controlling the false discovery rate: a practical and powerful approach to multiple testing. *Journal of the Royal statistical society: series B (Methodological)*, 57(1):289–300, 1995.

[11] Zachary D Kurtz, Christian L Müller, Emily R Miraldi, Dan R Littman, Martin J Blaser, and Richard A Bonneau. Sparse and compositionally robust inference of microbial ecological networks. *PLoS computational biology*, 11(5):e1004226, 2015.

[12] Grace Yoon, Irina Gaynanova, and Christian L Müller. Microbial networks in spring-semi-parametric rank-based correlation and partial correlation estimation for quantitative microbiome data. *Frontiers in genetics*, 10:449195, 2019.

[13] Nicolai Meinshausen and Peter Bühlmann. High-dimensional graphs and variable selection with the Lasso. *The Annals of Statistics*, 34(3):1436 – 1462, 2006.

[14] Jerome Friedman, Trevor Hastie, and Robert Tibshirani. Sparse inverse covariance estimation with the graphical lasso. *Biostatistics*, 9(3):432–441, 2008.
